# Lifestyle-specific responses of *Trichoderma* spp. in mycoparasitic confrontations and implications for biocontrol of *Populus* x *canescens*

**DOI:** 10.1101/2024.04.17.589723

**Authors:** Pia Stange, Johannes Kersting, Prasath Balaji Sivaprakasam Padmanaban, Jörg-Peter Schnitzler, Maaria Rosenkranz, Tanja Karl, J. Philipp Benz

## Abstract

- *Trichoderma* spp. are gaining popularity in agriculture and forestry due to their multifaceted roles in promoting plant growth through e.g. nutrient translocation, hormone production, induction of plant systemic resistance, but also direct antagonism of other fungi. However, the mycotrophic nature of the genus bears the risk of possible interference with other native plant-beneficial fungi, such as ectomycorrhiza, in the rhizosphere. Such interference could yield unpredictable consequences for the host plants of these ecosystems.
- We investigated whether *Trichoderma* spp. can differentiate between beneficial ectomycorrhizal fungi (represented by *Laccaria bicolor* and *Hebeloma cylindrosporum*) and pathogenic fungi (represented by *Fusarium graminearum* and *Alternaria alternata*) in different confrontation scenarios, including a newly developed olfactometer “race tube”-like system.
- Using two independent species, *T. harzianum,* and *T. atrobrunneum*, with plant-growth-promoting and immune-stimulating properties towards *Populus* x *canescens*, our study revealed robustly accelerated growth towards phytopathogens, while showing a contrary response to ectomycorrhizal fungi. Transcriptomic analyses identified distinct genetic programs during interaction corresponding to the lifestyles, emphasizing the expression of mycoparasitism-related genes only in the presence of phytopathogens.
- The findings reveal a critical mode of fungal community interactions belowground and suggest that *Trichoderma* spp. can distinguish between fungal partners of different lifestyles already at a distance.

## 1 Introduction

*Trichoderma* species, belonging to the phylum Ascomycota, are naturally occurring in soil as well as on plant surfaces (Afzal et al., 2021; Oskiera et al., 2017). The genus of *Trichoderma* has been widely studied as a biocontrol agent (BCA) (Bonfante & Genre, 2010; Guzmán-Guzmán et al., 2019; Sharma et al., 2009; Sood et al., 2020; Stenberg et al., 2021). Biocontrol is based on several mechanisms, such as the antagonistic activity against fungal plant pathogens, the proficiency to colonize plant tissues, the ability to induce systemic resistance in plants, as well as the adaptability to a wide range of environments (Liu et al., 2022; Manzar et al., 2022; Rush et al., 2021). During the complex process of mycoparasitism, coherent mechanisms of antibiosis, competition for nutrients and space and direct inhibition by the release of fungal cell wall degrading enzymes (CWDE), such as chitinases, proteases, and β-glucanases, can effectively lead to inhibition and death of prey fungi (Manzar et al., 2022; Sharma et al., 2011; Thambugala et al., 2020; Viterbo & Horwitz, 2010). The application of *Trichoderma* as BCA in disease management has already been shown to be effective against a broad range of foliar and root pathogens (Alfiky & Weisskopf, 2021; Benítez et al., 2005; Tyśkiewicz et al., 2022). As a result, more than 60 % of the officially registered BCAs are based on different species of *Trichoderma,* with *Trichoderma viride*, *Trichoderma virens,* and *Trichoderma harzianum* being the most commonly used (Abbey et al., 2019; Nur & Noor, 2020; Rush et al., 2021).

In addition to functioning as a bio-fungicide, numerous rhizosphere-competent *Trichoderma* spp. can form a close symbiosis with plant roots, producing soluble metabolites and volatile organic compounds (VOCs) with plant-performance stimulating activities conferring improved plant growth and induced resistance to abiotic stresses (Garnica-Vergara et al., 2016; Harman et al., 2004; Nawrocka & Małolepsza, 2013; Uniyal et al., 2018). Therefore, the application of *Trichoderma* spp. as an environmentally friendly bio-fertilizer and BCA can minimize the amount of traditional fertilizers by improving nutrient and water acquisition, as well as reducing the amount of synthetic fungicides (Hermosa et al., 2013; Hyakumachi & Kubota, 2004; López-Bucio et al., 2015; Shoresh et al., 2010). Moreover, the described beneficial effects imply the potential economic impact through shortened plant production cycles and increased germination rates of seeds in nurseries (Aleandri et al., 2015; Grodnitskaya & Sorokin, 2006).

Hybrid poplars (*Populus* spp.) have gained significance due to their fast growth, making them a valuable feedstock for a wide range of wood and non-wood products with high economic importance, including bioenergy, building materials, and paper production (Polle & Douglas, 2010; Przybysz & Przybysz, 2013; Rubin, 2008). *Populus* spp. are naturally found in complex symbiotic interactions with beneficial microbes such as ectomycorrhizal fungi (ECM) (Karliński et al., 2010; Kwaśna et al., 2021; Luo et al., 2009; Plett & Martin, 2012; Schnitzler et al., 2010). However, monocultures of poplar hybrids in large-scale short rotation coppices (SRC) are also often susceptible to a wide range of soil-born and foliar fungal pathogens such as *Armillaria* root rot, *Melampsora* leaf rust, or leaf spot caused by *Alternaria alternata* (Hacquard et al., 2011; Ostry et al., 2014). Despite *A. alternata* being recognized as an airborn pathogen, its chlamydospores have been observed to persist both in soil and infested organic matter and was found on diseased roots of *Vaccinium corymbosum* and *Taxus x media* (Nadziakiewicz et al., 2018). *Trichoderma* spp. isolated from the rhizosphere of various trees have been shown to display a high antagonistic activity against common poplar pathogens, including *A. alternata* (Asef et al., 2008; Yu et al., 2022). However, the mycotrophic nature of *Trichoderma* bears the potential to have also negative effects on the local ECM population through competition for essential nutrients or growth inhibition by direct antagonism. While *Trichoderma*-based BCA might thus be a promising tool in SRCs to mitigate susceptibility to phytopathogenic fungi and improve biomass productivity, further research is needed to study the interaction between *Trichoderma* and plant-beneficial fungi in the rhizosphere, such as ECM partners, to evaluate potential risks of adverse effects on these non-target associations (Minchin et al., 2012), as these interactions play crucial roles in soil health and nutrient uptake (Frąc et al., 2018; Ma et al., 2008; Mayor et al., 2015).

Existing literature reflects a degree of ambiguity regarding the ability of *Trichoderma* to differentiate between diverse fungal taxa, with important implications on the framework of the plant holobiont theory (Hassani et al., 2018; Zilber-Rosenberg & Rosenberg, 2008). Understanding whether *Trichoderma* can selectively interact with certain fungi while resisting/ignoring others would provide invaluable insights into the complex dynamics of plant-fungus interactions and ecosystem functioning. We therefore initiated a study to directly investigate the ability of *Trichoderma* spp. to distinguish between plant-beneficial ECM and plant-pathogenic fungi using representative species of each of these two major lifestyle groups. For this purpose, we selected two *Trichoderma* wild-type strains from the Harzianum clade (one *T. harzianum* and one *T. atrobrunneum* strain), which we tested for their biofertilizer and biocontrol capacity in grey poplar (*Populus* x *canescens*, syn. *P. alba* x *P. tremula*). Confrontation scenarios were set up with the phytopathogens *A. alternata* and *Fusarium graminearum* in direct comparison with *Laccaria bicolor* and *Hebeloma cylindrosporum* as representative plant-mutualistic ECM. Moreover, to evaluate the physiological response of *Trichoderma* over longer distances and time while having a two-directional choice of growth, we developed a novel olfactometer “race tube”-like system. Transcriptomic analysis was used to detect differences in the initiation of mycoparasitism-related programs during the different contact stages and better understand the interaction on a molecular level.

## 2 Materials and Methods

### Cultures and growth conditions

The *Trichoderma* strain *T. harzianum* WM24a1 was obtained from the Austrian Institute of Technology GmbH (Monika Schmoll; Tulln, Austria), and *T. atrobrunneum* was isolated from a wood sample in 2018 (Bavaria, Germany). Both strains were identified previously on a molecular level following Cai & Druzhinina (2021) and tested *in vitro* for their biocontrol capacities (Stange et al., 2023). As potential preys the ECM basidiomycetes *Laccaria bicolor* isolate S238N (Institute National de la Recherche Agronomique, Nancy, France) and *Hebeloma cylindrosporum* (obtained from Technical University Dresden, Germany) and the plant pathogens *Fusarium graminearum* PH-1 (obtained from University of Hamburg, Germany) and *Alternaria alternata* 22-2 (Phytopathology, Technical University of Munich) were used. Agar plugs from ECM fungi and *F. graminearum,* and spores from *Trichoderma* and *A. alternata* were routinely sub-cultured on potato dextrose agar and incubated at 21 °C and 75 % humidity in constant darkness. *Populus* x *canescens* INRA clone 717 1-B4 was micropropagated routinely in Schenk and Hildebrandt medium (SH medium, Schenk & Hildebrandt, 1972) as described by Behnke et al., 2007 and Müller, Volmer et al., 2013 and cultivated at 21°C, 75 % humidity, and 16 h photoperiod with 105 µmol^-2^s^-1^ (daylight white color 865).

### Biofertilizer and biocontrol capacity in poplar

Test plants (Supplementary Methods **S1**) were inoculated with five ml spore suspension of *T. harzianum* WM24a1 and *T. atrobrunneum* containing 10^6^ spores ml^-1^ in sterile water, control plants were mock inoculated with sterile water. Directly after transfer and every two weeks plants were watered with ten ml of ¼ strength Long Ashton nutrient solution (Hewitt & Smith, 1975). The 5-week-old plants were inoculated again with five ml of a spore suspension containing 1 x 10^6^ spores ml^-1^. After four days, one leaf per plant was wounded at four sites with a sterile needle. For infection with *A. alternata* five µl of a spore solution containing 3 x 10^6^ spores ml^-1^ was pipetted directly to the wounding site. For mock inoculation autoclaved water without spores was used and five replicates for each treatment were prepared. Pictures of the leaves were taken after five days and the total infection area per leaf was measured using ImageJ V1.53e software (Wayne Rasby, National Institute of Health, USA, http://imageJ.nih.gov/ij). Plant height, leaf number and shoot and root fresh weight were assessed for a subset of plants (*n* = 7) after 6 weeks. Dry weight was determined after 48 h at 50°C.

### Antagonistic activity of *Trichoderma*

For *in vitro* antagonism assays in dual culture, the second fungal partners *H. cylindrosporum*, *L. bicolor*, *F. graminearum,* or *A. alternata* were inoculated on solid Modified Melin-Norkrans synthetic medium (MMN) (Müller; Volmer et al. 2013) in non-split and split Petri dishes (9 cm diameter) (Guo et al., 2019) and the ECM were incubated for two and the pathogens one weeks, respectively. *Trichoderma* strains were inoculated on the other side of the plate as spores and after three days photos were taken with a Nikon camera (Nikon, Tokyo, Japan) The colony area (cm^2^) of fungal mycelium in media contact (MC) and air contact (AC) was measured using ImageJ. As a control, each fungus was grown alone. The inhibitory effect was calculated by using the following formula (Raut et al., 2014):

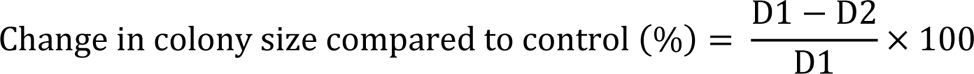

With D1 = colony area of control condition and D2 = colony area of test condition

### Physiological response of *Trichoderma* using race tube system

The olfactometer “race tube”-like system (Supplementary Fig. **S2,** Methods **S2**) was composed of two 220 ml sample cups (391-0023, VWR, Darmstadt, Germany) and a 50 ml serological pipette (612-3696, VWR, Darmstadt, Germany). *L. bicolor* and *H. cylindrosporum* as well as *F. graminearum* and *A. alternata* were inoculated into the right tube two and one week before *Trichoderma* was inoculated with spore solution into the middle of the race tube. For AC condition the second fungus was inoculated into a small petri dish and placed into the test cup. Both cups were sealed with a gas-permeable membrane (BR701364, Sigma, St. Louis, USA). The left tube remained uninoculated, serving as a control tube. Growth of *Trichoderma* was monitored at 48 h, 72 h, 96 h, 120 h, and 144 h, respectively, after inoculation by marking the hyphal growth on the corresponding section of the serological pipette. As a control, *Trichoderma* was challenged with itself. The direction of hyphal growth of *Trichoderma* was determined by the percentage of total growth between the described time points and Δ % of growth was calculated with the following formula:

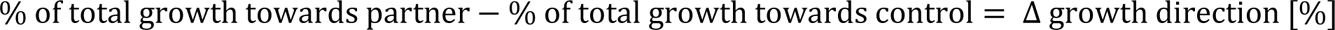

### Transcriptomic analysis of confrontations with phytopathogens and ectomycorrhiza

To determine whether the presence of a plant-beneficial and a plant-pathogenic prey affects the expression of mycoparasitism-related genes during different degrees of contact with ECM and plant-pathogens, an RNA-Seq experiment was performed. Confrontation scenarios where the same as for testing antagonistic activity, consistent with modified MMN media overlaid with wet autoclaved cellophane membrane (Natureflex 32g m^-2,^ HERA, Schotten, Germany) to enable biomass harvest without agar residues. The biomass was harvested three and six days after *Trichoderma* inoculation with a sterile spatula and subjected to RNA extraction (Supplementary Methods **S3**). Qualified RNA was subjected to cDNA library preparation using the Illumina stranded mRNA-Kit (Illumina, San Diego, USA) quantified and qualified (DNA High Sensitivity assay, Aligent Technologies, Santa Clara, USA) by the chair of Animal Physiology and Immunology at TUM. Sequencing of barcoded libraries was done at IMGM laboratories (Martinsried, Germany) with NovaSeq 6000 for 100 bp single-end reads to a depth of 18 million reads per sample. *H. cylindrosporum* could not be included in this transcriptomic analysis, since it was not available at that time.

### Bioinformatic analysis

Sequencing data was processed using nf-core/rna-seq v3.12.0 workflow (Ewels et al., 2020) and executed with Nextflow v23.10.0 (Di Tommaso et al., 2017). The reads were trimmed using Cutadapt (v4.8) (Martin, 2011) to remove poly-A tails, adaptor sequence contaminations and low-quality bases and subsequently aligned to the concatenated reference genomes with STAR (Dobin et al., 2013). Gene-level read counts were determined using Salmon (Patro et al., 2017) and subjected to downstream analysis. Differential gene expression analysis was conducted using the DESeq2 R package (v1.16.1) to normalize the libraries based on the geometric mean of the read counts and then calculate the log2fold change (LFC) between the experimental test conditions and the control condition (Love et al., 2014). Genes were identified to be differentially expressed (DEGs) with adjusted *p*-value (FDR) < 0.01. Upregulated DEGs (LFC > 0) were submitted to FungiFun 2.2.8 (Priebe et al., 2015) with *T. harzianum* CBS 226.95 and *L. bicolor* S238N-H82 / ATCC MYA-4686 as reference species, to assign functional annotations of DEGs and conduct enrichment analyses. DEGs were classified based on Functional Catalogue (FunCat) (Ruepp et al., 2004), Gene Ontology (GO) (Harris et al., 2004), and Kyoto Encyclopedia of Genes and Genomes (KEGG) (Kanehisa & Goto, 2000) and tested for enrichment using FungiFun 2.2.8 BETA (Priebe et al., 2015). The presence of signal peptides in DEGs was predicted with SignalP 5.0 (Almagro Armenteros et al., 2019). Self-organizing tree algorithm (SOTA) was used to cluster common DEGs based on expression patterns between the different test conditions using Pearson’s correlation and MeV v4.4.1 (Howe et al., 2011).

### Statistical analysis and visualization

The obtained data was processed for statistical data analysis with Origin Pro (version 2021, OriginLab Corporation, Northampton, USA). Data was tested for normal distribution using the Shapiro-Wilk test, followed by Levene’s test for variance homogeneity. One-way ANOVA was performed combined with Bonferroni test for pairwise comparison. For plant data, one-way ANOVA was applied combined with Fisher’s least significant difference (LSD) test. Figures were created and combined with BioRender.com (TUM institutional license WN26OFV8MS).

## 3 Results

### *Trichoderma* spp. are displaying bio-fertilizer and biocontrol capacity in poplar

When considering *Trichoderma* strains as a BCA for poplar plantations, the potential to induce effective plant systemic resistance may vary between different strains used. It is therefore important to exclude potential negative effects by evaluating the interaction of the biocontrol strain and the plant. The biofertilizer capacity was evaluated by comparing plant height, leaf number, shoot and root fresh and dry weight to the untreated control plants (Fig. **1**). The chosen strains significantly increased plant height, average leaf number (from 7.2 (+/- 1.6) to 8.4 (+/- 0.9) with *T. harzianum* treatment and 9.4 (+/- 1.2) for *T. atrobrunneum* treatment) (Fig. **1a,b**), as well as shoot and root fresh weight (Fig. **1c,d**). To evaluate the potential biocontrol capacity of both strains, one leaf of each poplar-plant was injured and infected with spores of *A. alternata*. Both *Trichoderma* strains significantly decreased signs of infection in leaves (Fig. **1g**) reflecting a positive influence on the induced systemic resistance in *P.* x *canescens*.

**Fig. 1:**
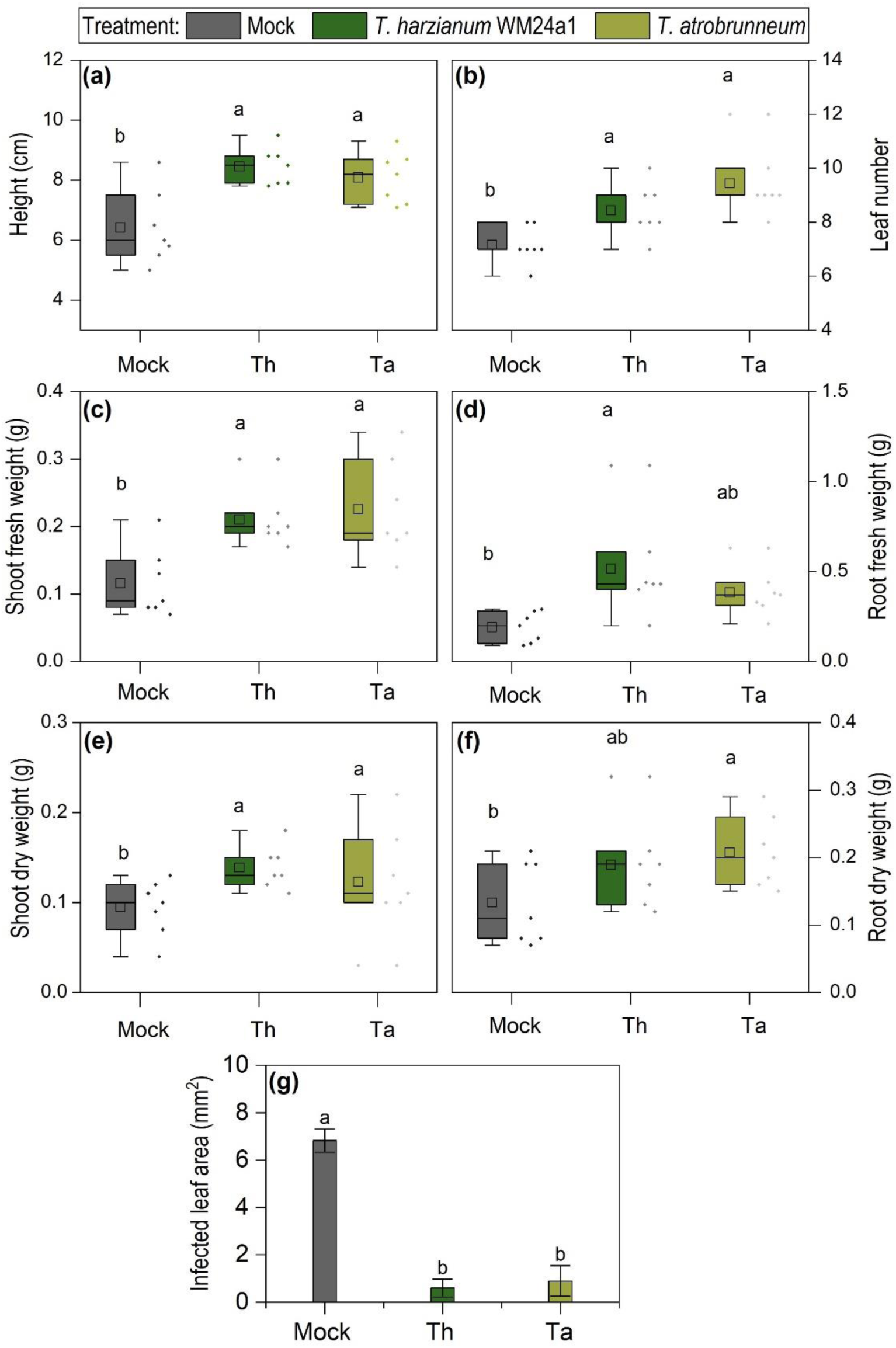
Evaluation of biofertilizer and biocontrol capacity of *T. harzianum* WM24a1 and *T. atrobrunneum* in *Populus x canescens*. Plant height **(a)**, leaf number **(b)**, shoot fresh and dry weight **(c,e)** and root fresh and dry weight **(d,f)** were assessed after six weeks of cultivation (*n* = 7). A subset of plants were inoculated again with 5 ml spore solutions containing 10^6^ spores ml^-1^. After 4 days one leaf per plant was injured with a sterile needle and infected with *A. alternata* spore solution (*n* = 5). Infection area in mm^2^ **(g)**. Significances were determined by one-way ANOVA and LSD, p < 0.05.

### Differences in antagonistic activity of *Trichoderma* towards plant-beneficial and plant-pathogenic fungi

To examine the physiological response and potential inhibitory effect of *Trichoderma* species towards plant-pathogenic and plant-beneficial fungi, we initially set up dual confrontation assays with *F. graminearum*, *A. alternata, L. bicolor* and *H. cylindrosporum*, in different degrees of contact in standard Petri dish systems (Supplementary Fig. **S1**).

A clear inhibitory effect of both *Trichoderma* strains towards all confrontation partners was observed three days after inoculation with *Trichoderma*, albeit differing in severity depending on the partner and being generally stronger during conditions that allowed media contact (MC), compared to the situation in a split-plate that allowed only contact through the headspace (air contact; AC). Also, *T. harzianum* showed a stronger inhibitory effect on both pathogens during MC compared to AC, whereas *T. atrobrunneum* only showed stronger inhibitory effects during MC towards *A. alternata* (Fig. **2c,d**). The inhibitory effect of *Trichoderma* towards the ECM was stronger towards *H. cylindrosporum* compared to *L. bicolor* for both strains (Fig. **2a,b**). On the contrary, the presence of *H. cylindrosporum* and *L. bicolor* was found to inhibit the growth of both *Trichoderma* strains (between 4% and 47%) (Fig. **2b**), and this effect was more pronounced, albeit not significantly, in the AC conditions and somewhat more robust for *L. bicolor* than for *H. cylindrosporum* (Fig. **2a,b**). At the same time, the presence of both pathogens led to an enhanced colony diameter of *Trichoderma*, indicated by negative inhibition values (Fig. **2c,d**). In *T. atrobrunneum*, this effect ranged between 2% and 10% and in *T. harzianum* between 11% and 29% compared to the *Trichoderma*-only control. Intriguingly, while this increase was stronger during AC confrontation with *F. graminearum* compared to MC confrontation, no significant differences could be detected between AC and MC confrontation with *A. alternata*. Overall, these data show that while the ECM fungi have a robust repelling effect on *Trichoderma* spp., they seem to be attracted to plant-pathogenic fungi.

**Fig. 2:**
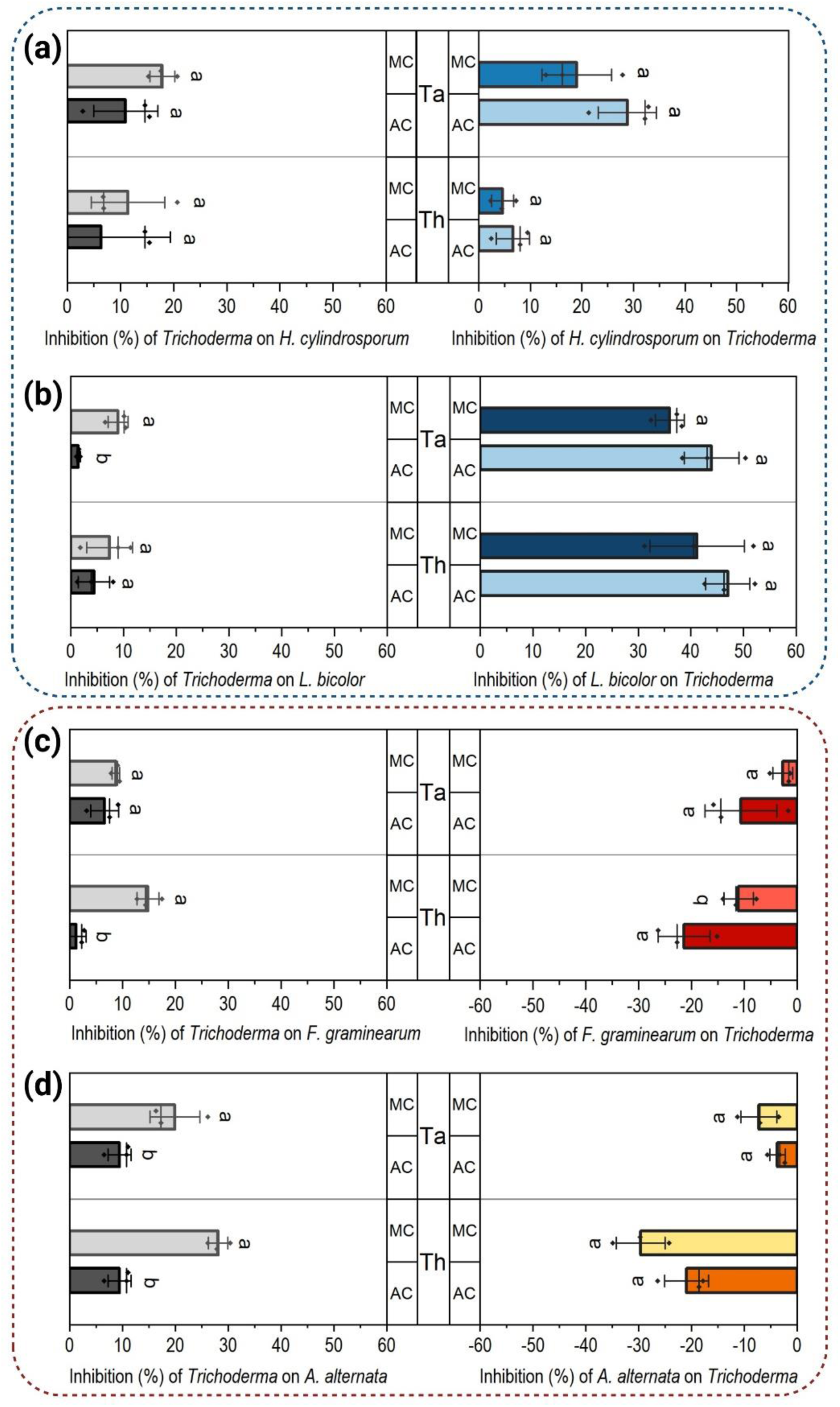
Growth inhibition during co-cultivation of *Trichoderma* spp. with the two ectomycorrhizal fungi *H. cylindrosporum* **(a)** and *L. bicolor* **(b)** and the two plant-pathogenic fungi *F. graminearum* **(c)** and *A. alternata* **(d)**. Growth inhibition of each fungus during co-cultivations was calculated by comparing the colony area with single controls for MC and AC three days after inoculation with *Trichoderma*. Significances were determined for each contact stage separately with Student’s *t*-test, p < 0.05, *n* = 3.

### A novel olfactometer “race tube”-like system to quantify the directional growth response to fungal partners

To overcome the limitations of more traditional plate confrontation assays, we developed a novel olfactometer “race-tube”-like system. Differing from the Petri dish system, this experimental set-up allows for a two-way choice of growth direction and therefore an observation of the directed growth of *Trichoderma* in presence of a second fungus (Fig. **3a**). Furthermore, the growth direction can be evaluated and quantified over much longer distances and time and therefore in a much more reliable fashion. Self-confrontation of both *Trichoderma* strains led to Δ growth direction values near 0, indicating an equal growth towards both directions (Fig. **3b-e**). Using the new system, we could confirm the overall effects observed in the plate-based system. However, it became obvious that both *Trichoderma* strains were not simply inhibited by the ECM fungi, but indeed chose to grow away from them, visible by stronger growth in the opposite direction, leading to negative Δ growth direction values. This effect was stronger for *T. harzianum* during AC compared to MC (Fig. **3b,c**), whereas it was the other way around for *T. atrobrunneum* (Fig. **3d**). The situation was found to be completely different in presence of phytopathogens, and both *Trichoderma* strains displayed a clear growth preference towards those fungi, as indicated by positive Δ growth direction values (Fig. **3b-e**). In *T. harzianum* this effect was stronger during the first 72 h in AC compared to MC, shifting to more pronounced effects in MC compared to AC after 96 h (Fig. **3b,c**). Interestingly, in both *Trichoderma* strains the directed growth towards the pathogens in AC confrontation decreased slightly after 96 h, and in MC confrontation after 120 h. Overall, the physiological responses indicated a negative chemotropism in presence of ECM and a positive chemotropism in presence of phytopathogenic fungi taking effect already at comparably long distances.

**Fig. 3:**
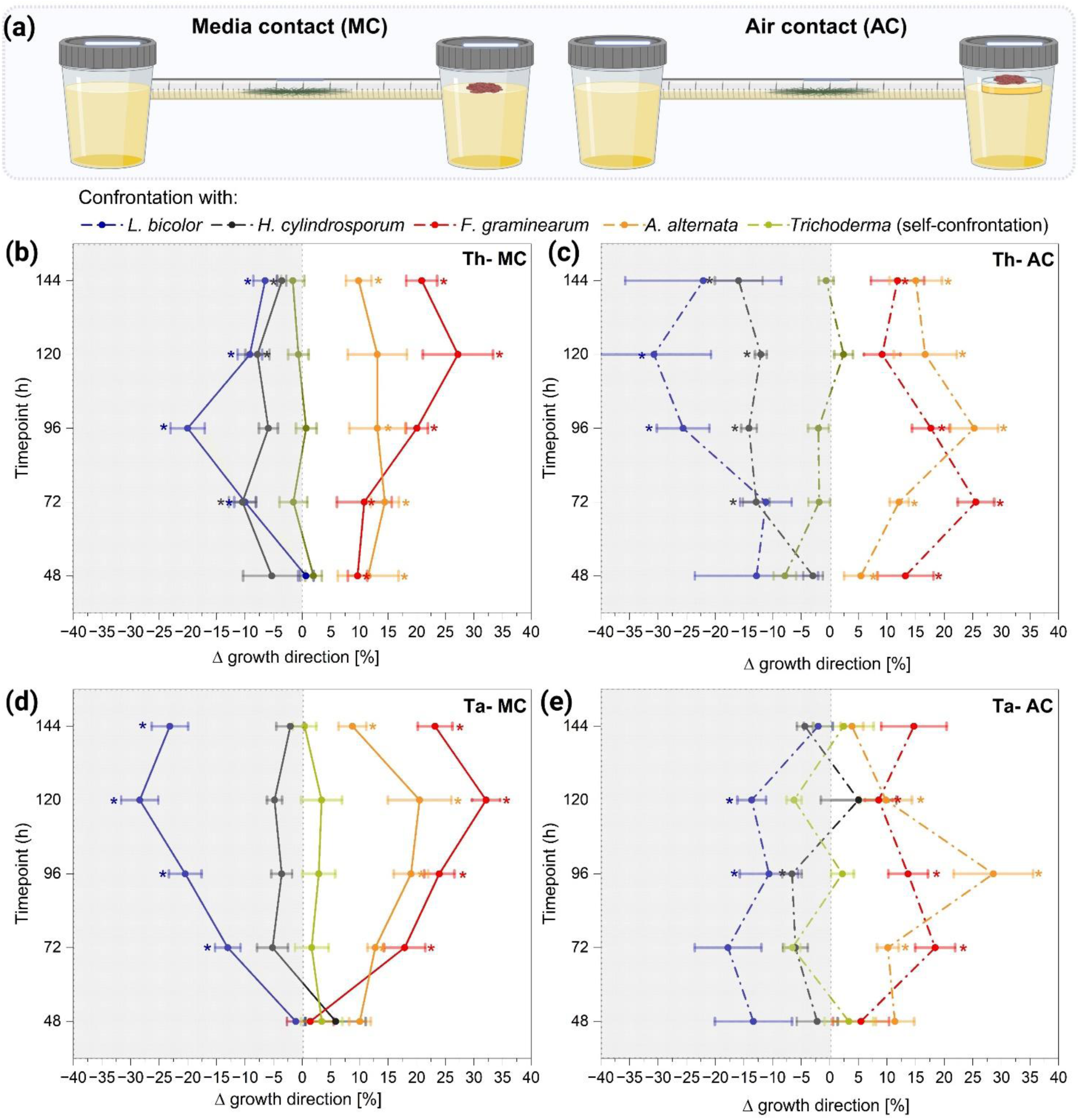
Experimental setup of olfactometer “race tube”-like system and the confrontations with fungal partner during media (MC) and air contact (AC) **(a)**. Physiological response of *T. harzianum* WM24a1 **(a,c)** and *T. atrobrunneum* **(d,e)** in the presence of *H. cylindrosporum* (grey), *L bicolor* (blue), *F. graminearum* (red) and *A. alternata* (orange). The direction of growth was determined as Δ growth direction (%) between the different time points. As a control, *Trichoderma* was challenged with itself (green). Significances were determined with Student’s *t*-test compared to the self-confrontation, p < 0.05, *n* = 5.

### Distinct patterns in *Trichoderma* global gene expression

To identify differentially expressed genes (DEGs) related to the strongly differing reaction of *Trichoderma* to ECM and phytopathogenic fungi, another series of plate-based confrontations was conducted with *T. harzianum* on one side and *L. bicolor*, *F. graminearum*, or *A. alternata* on the other. The plate system was used in this experiment, since it allowed harvesting biomass directly from the zone of interaction (Fig. **4a**). Samples were taken three days after inoculation before any kind of physical contact (MC) and six days after inoculation when a direct hyphal interaction was established (DC).

**Fig. 4:**
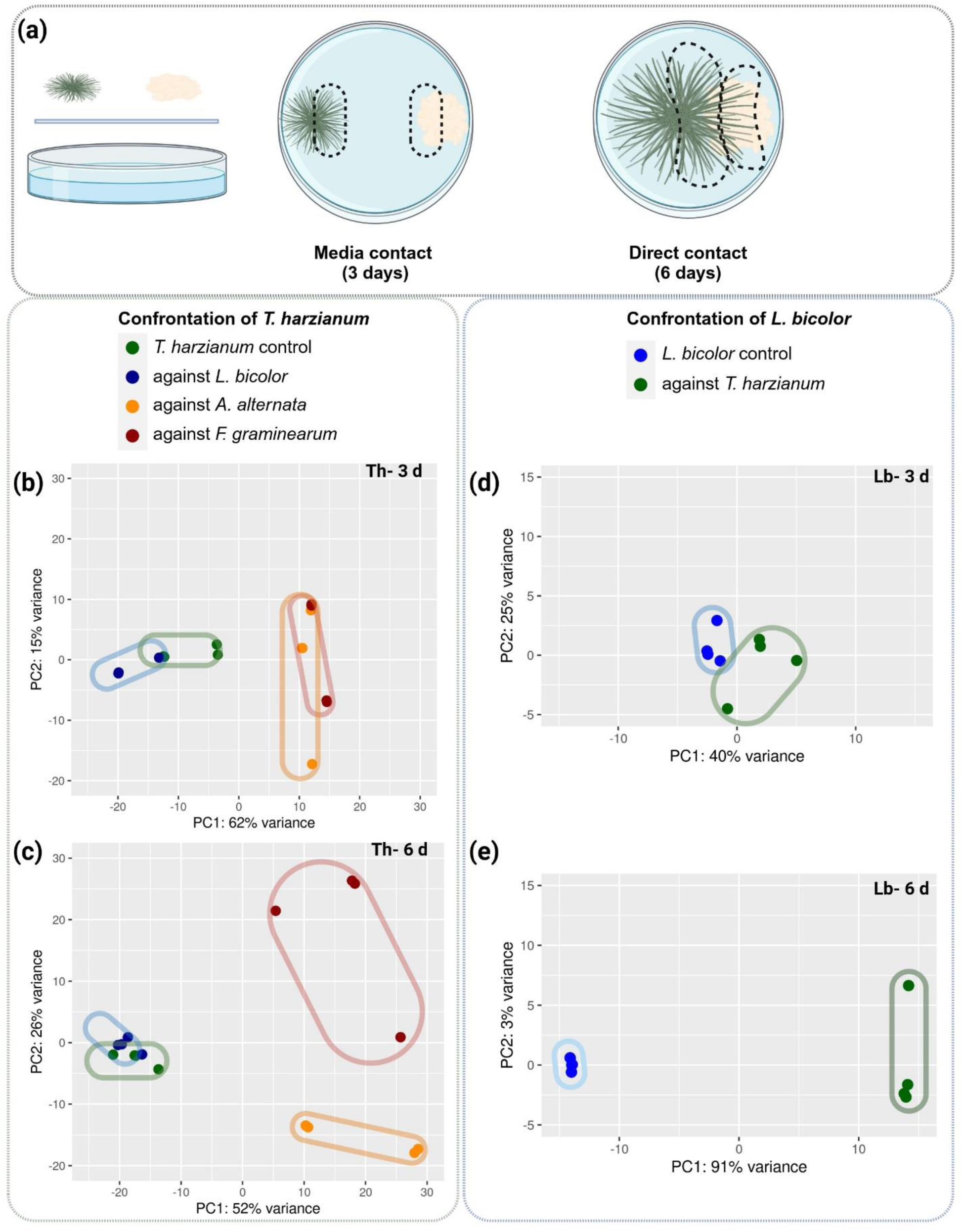
Transcriptomic analysis of fungal confrontations. Experimental setup alongside with depicting areas of biomass harvest from the zone of interaction **(a)**. PCA plots illustrate differential gene expression in *T. harzianum* after three (MC) and six days (DC) **(b,c)** and in *L. bicolor* **(d,e)** based on the top 500 differentially expressed genes among all conditions.

Compared to the concatenated reference genomes of *T. harzianum* CBS 226.95 and *Laccaria bicolor* S238N-H82, the alignment rate ranged from 81.2% to 97.7%, respectively. Principal component analysis (PCA) revealed cluster formation of the confrontations with the pathogens away from the *Trichoderma*-only control, indicating similarly changed expression patterns, while confrontation with *L. bicolor* clustered much closer to the controls, indicating more limited changes (Fig. **4b**). At DC, the pathogen interactions separated into individual clusters, while the confrontation with *L. bicolor* still clustered with the control (Fig. **4c**). This differential clustering highlights the similarities between confrontation with *L. bicolor* and the control condition and points to distinct expression patterns during the confrontation with the pathogens developing after three days.

Distinct patterns across the confrontations with the ECM and the pathogens during MC emerged also when looking at the gene expression more closely (volcano plots; Supplementary Fig. **S3**). The confrontation with *L. bicolor* was characterized by small LFC values compared to the control, representing minimal changes in gene expression. To nevertheless ensure not to lose potentially relevant genes in the downstream analysis, we employed a threshold of adjusted FDR < 0.01 without an additional LFC threshold. Especially during the MC stage without physical contact, we expected the potential signaling mechanisms not to display extreme fold-changes, which should nevertheless be significant. This approach allowed to capture the nuanced variations in gene expression during confrontation with the ECM. Conversely, in confrontation with the pathogens during the MC, the volcano plots illustrate a broader dispersion of points with higher LFC values, indicating a notable and pronounced alteration in gene expression.

### Unique genetic response of *T. harzianum* towards *L. bicolor*

Overall, the interaction of *T. harzianum* and *L. bicolor* during MC (three days) led to the up- and downregulation of only 67 and 56 DEGs in *T. harzianum*, respectively (Fig. **5a**). This observation aligns with the previous observations. The changes in the transcriptome of *T. harzianum* were small compared to the distinct and pronounced changes during confrontation with *F. graminearum* and *A. alternata*, which led to 614 and 1,072 upregulated DEGs and 1,262 and 893 downregulated DEGs, respectively. During DC (day six) 366 and 175 genes were significantly up- and downregulated in *T. harzianum* during interaction with *L. bicolor*, whereas the presence of both pathogens led to much more DEGs (2,366 and 1,692 upregulated DEGs during confrontation with *A. alternata* and *F. graminearum*, respectively) (Fig. **5a**). After three days of confrontation with the two pathogens, 767 and 483 DEGs were commonly up- and downregulated in *T. harzianum* (Fig. **5c,d**), while during confrontation with *L. bicolor* (TL3), 65 and 46 were uniquely up- and downregulated, respectively. During overgrowth stage (DC, six days), confrontation with pathogens up- and downregulated 873 and 831 shared DEGs in *T. harzianum*, while only 75 and 58 DEGs were common also in the confrontation with *L. bicolor* (Fig. **5e,f**).

**Fig. 5:**
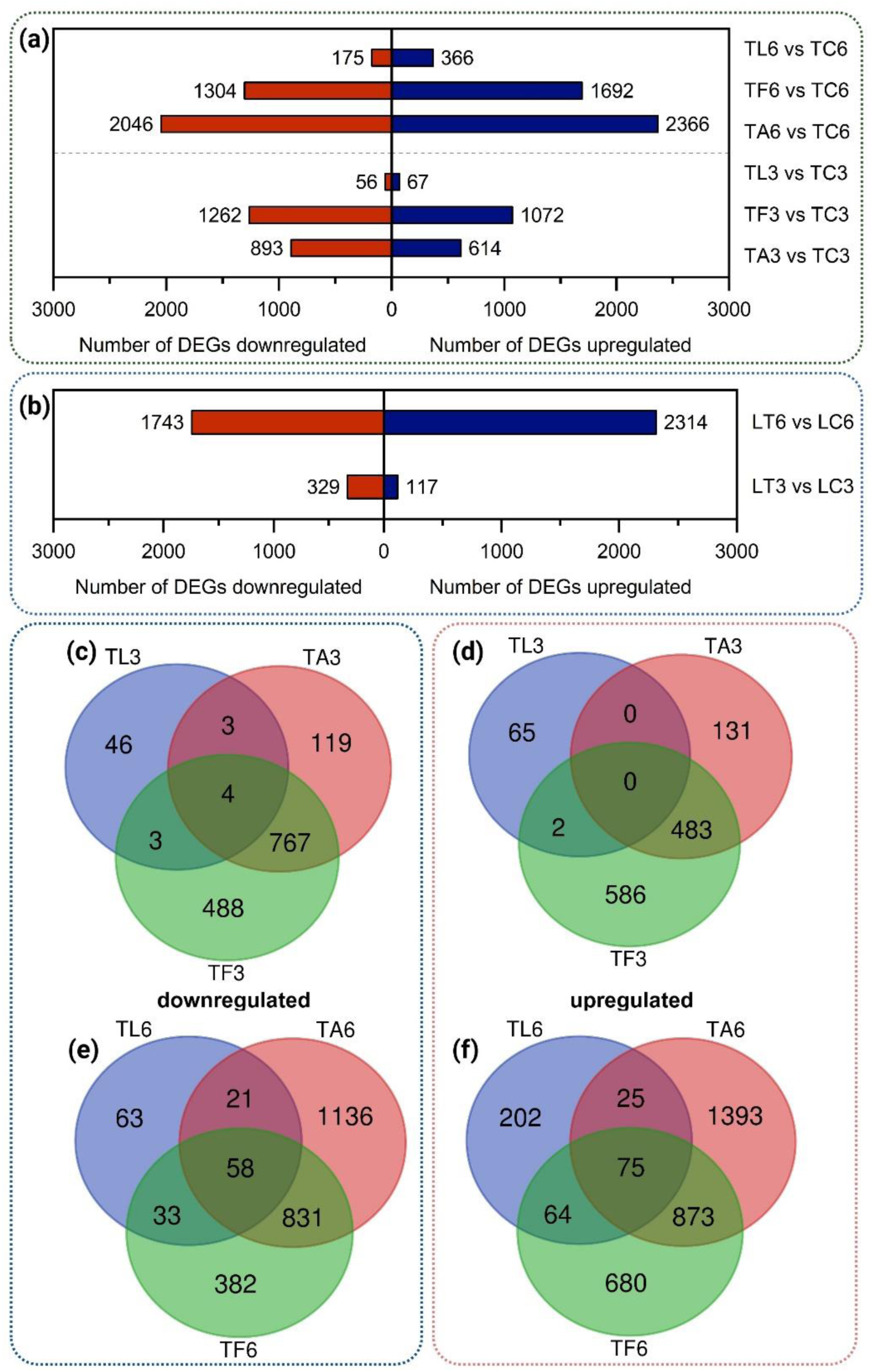
Number of DEGs (FDR < 0.01) that were up- or downregulated in the confrontation of *T. harzianum* with *L. bicolor* (TL), *A. alternata* (TA), and *F. graminearum* (TF) compared to the control condition after three days (MC) (TL3, TA3, TF3) and six days (DC) (TL6, TA6, TF6) **(a)**. Number of DEGs (FDR < 0.01) that were up- or downregulated in the confrontation of *L. bicolor* with *T. harzianum* (LT) after three and six days (LT3 and LT6, respectively) **(b)**. Venn diagrams showing common and unique DEGs among the three different confrontations of *T. harzianum* with *L. bicolor* (TL), *A. alternata* (TA), and *F. graminearum* (TF) after three days **(c,d)** and after six days **(e,f)**.

During MC (three days), at least 13 genes from among the common and most upregulated 30 DEGs in confrontation with *A. alternata* (TA3) and *F. graminearum* (TF3) are known to play a crucial role in the process of mycoparasitism, clearly showing induction of related gene cascades already long before direct hyphal contact. On the contrary, during confrontation with *L. bicolor* (TL3) those mycoparasitism-related genes rather showed a downregulation (Table **1**). Interestingly, a terpene synthase (M431DRAFT_113113; LFC −0.9) was significantly downregulated as well. Furthermore, two short and uncharacterized signal peptide-containing proteins (M431DRAFT_69921, M431DRAFT_129453) were significantly induced.

**Table 1:** Genes in *T. harzianum* with known or predicted functions in the process of mycoparasitism that were found to be upregulated during the “sensing-phase” in presence of *A. alternata* (TA3) and *F. graminearum* (TF3), but not *L. bicolor* (TL3), at 3 days post inoculation. Predicted protein names were retrieved from UniProt and log2 fold-changes (log2FC) of all three confrontations are displayed and highlighted red for log2FC < 0, blue 0 > log2FC > 1 and green for log2FC > 1. Genes which were removed due to low counts show log2FC of NA. DEGs containing a signal-peptide predicted by SignalP are marked with a **^†^**.

| Predicted protein name | Gene ID | log2FC TA3 | log2FC TF3 | log2FC TL3 | Assigned GO terms |
| --- | --- | --- | --- | --- | --- |
| glucan endo-1,3-beta-D-glucosidase† | M431DRAFT_479664 | 2.84 | 4.50 | 0.13 | beta-glucosidase activity [GO:0008422]; polysaccharide catabolic process [GO:0000272] |
| Carbohydrate-binding module family 24 protein† | M431DRAFT_525334 | 1.87 | 3.56 | - 0.53 | glucan endo-1,3-alpha-glucosidase activity [GO:0051118] |
| Oligopeptide transporter | M431DRAFT_529850 | 2.45 | 3.51 | - 0.67 | oligopeptide transmembrane transporter activity [GO:0035673]; membrane [GO:0016020] |
| Peptidase S1 domain-containing protein† | M431DRAFT_526221 | 2.13 | 3.37 | - 0.62 | serine-type endopeptidase activity [GO:0004252]; proteolysis [GO:0006508] |
| Glycoside hydrolase family 18 protein† | M431DRAFT_509593 | 2.48 | 3.30 | - 0.23 | hydrolase activity [GO:0016787]; carbohydrate metabolic process [GO:0005975] |
| Carbohydrate-binding module family 13 protein | M431DRAFT_9084 | 1.73 | 2.17 | - 0.35 | carbohydrate metabolic process [GO:0005975] |
| Glycoside hydrolase family 3 protein, β-glucosidase<sup>†</sup> | M431DRAFT_504078 | 1.65 | 1.97 | - 0.51 | hydrolase activity, hydrolyzing O-glycosyl compounds [GO:0004553]; polysaccharide catabolic process [GO:0000272] |
| Chitinase<sup>†</sup> | M431DRAFT_505895 | 1.46 | 1.95 | - 0.32 | chitin binding [GO:0008061]; chitinase activity [GO:0004568]; chitin catabolic process [GO:0006032]; polysaccharide catabolic process [GO:0000272] |
| Chitinase<sup>†</sup> | M431DRAFT_500888 | 2.30 | 1.89 | NA | cellulose binding [GO:0030248]; chitinase activity [GO:0004568]; chitin catabolic process [GO:0006032]; polysaccharide catabolic process [GO:0000272] |
| Amino acid permease/ SLC12A domain-containing protein | M431DRAFT_92558 | 1.07 | 1.55 | - 0.44 | membrane [GO:0016020]; amino acid transport [GO:0006865]; transmembrane transport [GO:0055085] |
| Major facilitator superfamily (MFS) profile domain-containing protein | M431DRAFT_101977 | 1.65 | 1.52 | 0.84 | transmembrane transporter activity [GO:0022857]; membrane [GO:0016020] |
| Glycosyl-transferase family 32 protein | M431DRAFT_238777 | 0.94 | 1.49 | - 0.17 | alpha-1,6-mannosyltransferase activity [GO:0000009]; carbohydrate derivative metabolic process [GO:1901135] |
| Chitinase<sup>†</sup> | M431DRAFT_517960 | 0.74 | 1.37 | 0.05 | chitin binding [GO:0008061]; chitinase activity [GO:0004568]; chitin catabolic process [GO:0006032]; polysaccharide catabolic process [GO:0000272] |

Gene ontology (GO) analysis of upregulated genes showed clear enrichment (exact Fisher test, adj. *p*-value > 0.05) of terms involved in primary metabolic activity, such as “ribosomes”, “translation” and “carbohydrate metabolic process” (Fig. **6a**). This aligns with the phenotypic observations of increased colony area when *T. harzianum* was confronted with plant pathogens. Furthermore, GO terms of “cellulose binding”, “chitinase activity” and “hydrolase activity” were significantly enriched, corroborating that *Trichoderma* is able to sense potential prey already before direct hyphal contact. Conversely, GO enrichment analysis of upregulated DEGs in interaction with *L. bicolor* revealed a much more limited outcome with membrane-related annotation being the only enriched category (Fig. **6b**).

**Fig. 6:**
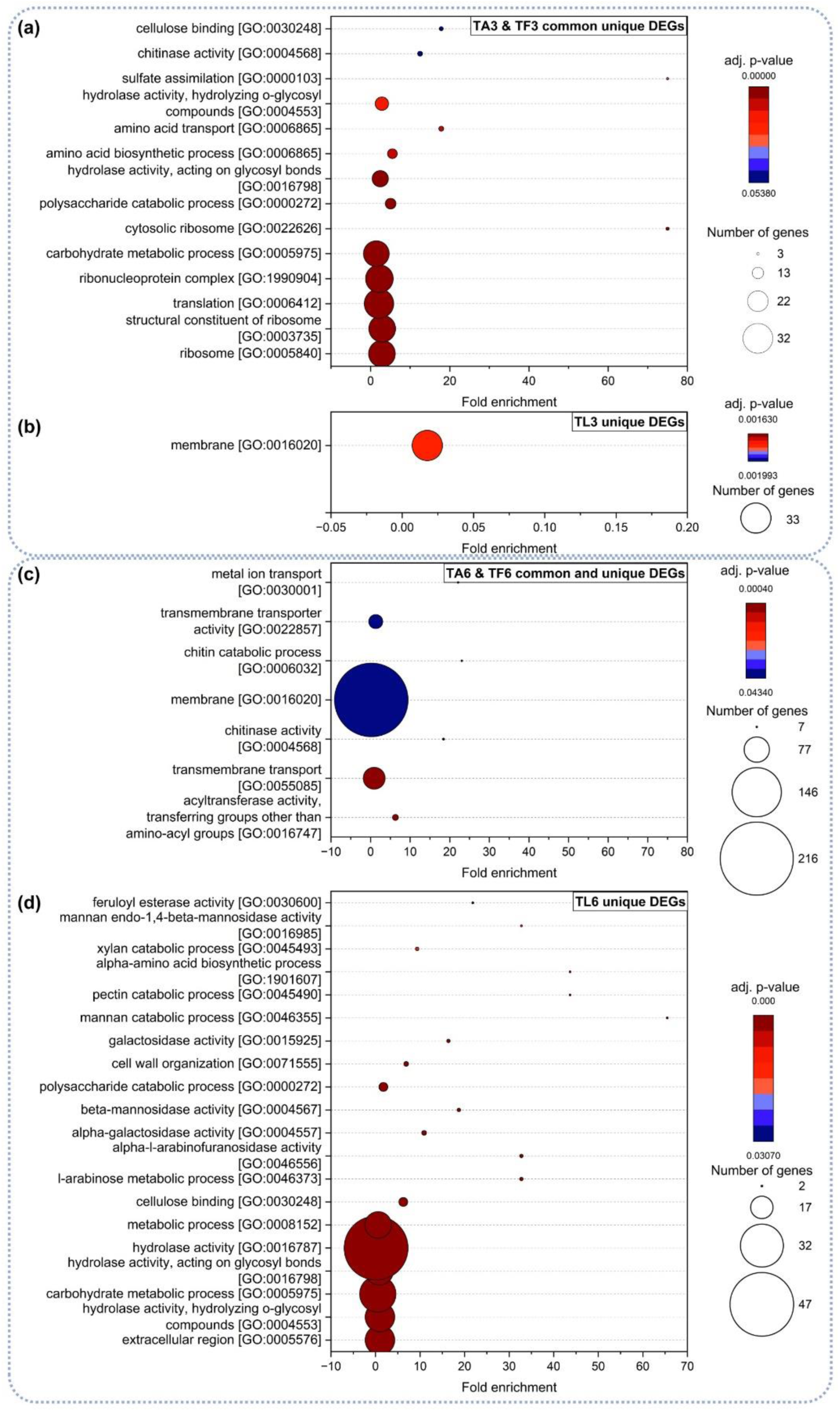
Gene ontology enrichment analysis of common and unique upregulated DEGs in *T. harzianum* during the interaction with *A. alternata* (TA3, TA6) and *F. graminearum* (TF3, TF6) after three days (MC) **(a)** and six days (DC) **(c)**. GO enrichment analysis of DEGs uniquely upregulated in *T. harzianum* in presence of *L. bicolor* after three days (TL3) **(b)** and six days (TL6) **(d)**. GO terms were assumed to be significantly enriched with adjusted *p*-value < 0.05.

Intriguingly, we identified 57 genes that were differentially expressed in all three test conditions during MC (three days), but which nevertheless show distinct expression patterns, according to the lifestyles. Expression profiles of those common DEGs by SOTA method using Pearson’s correlation revealed two distinct clusters (Fig. **7**, Supplementary Fig. **S4,** Table **S2**): 42 genes (cluster 1) showed a significant induction in *T. harzianum* during confrontation with *L. bicolor* and significant repression during confrontation with the pathogens. The remaining 15 DEGs (cluster 2), showed an opposite regulation, being significantly upregulated during the confrontation with the pathogens and downregulated during confrontation with *L. bicolor*. Several DEGs in cluster 1 are assigned to GO terms of “membrane” and associated with transport activities, as seen above. Interestingly, also a small secreted protein (M431DRAFT_96469; signal peptide likelihood: 0.99) was significantly induced in confrontation with *L. bicolor* with an LFC of 0.83 (FDR 0.001) and downregulated in presence of *A. alternata* and *F. graminearum* with LFC of −0.99 and −1.15 (FDR 2.75E^-8^ and 1.45E^-6^), respectively. Cluster 2 contains several DEGs annotated with GO terms of “hydrolase activity” and a peptidase A4 family protein upregulated with LFCs of 0.80 and 1.01 (FDR 2.96E^-11^ and 1.23E^-7^) in confrontation with *A. alternata* and *F. graminearum*, respectively, and downregulated in confrontation with *L. bicolor* (LFC of −1.19; FDR 3.7E^-14^).

**Fig. 7:**
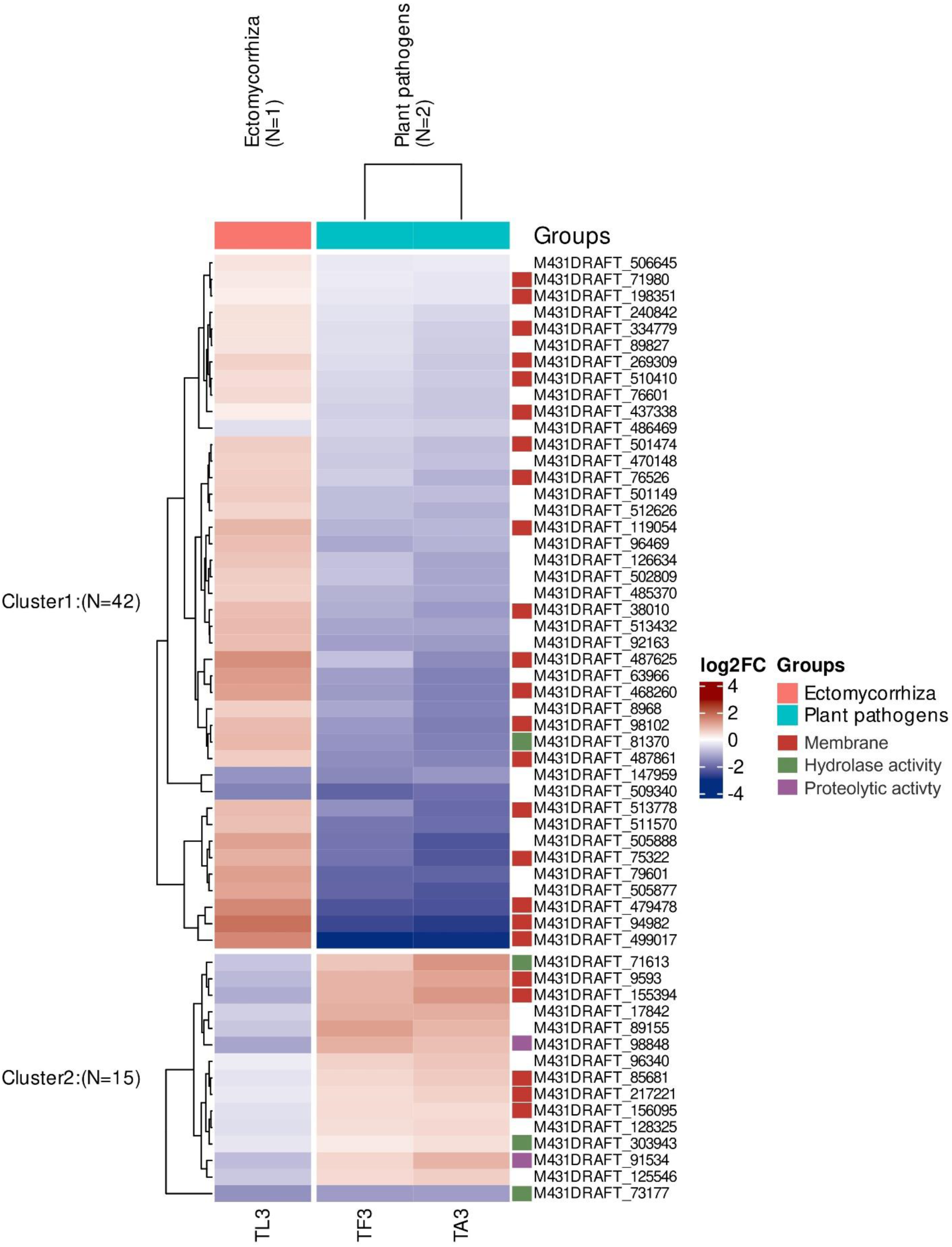
Heatmap of the 57 common DEGs in *T. harzianum* after three days during confrontation with *L. bicolor* (TL3), *A. alternata* (TA3) and *F. graminearum* (TF3) shown as log2 fold change (normalized to single control of *T. harzianum* TC3) and clustered based on expression using SOTA.

GO-term enrichment analysis of *T. harzianum* genes upregulated at day six of contact with both pathogens identified “membrane”-related categories, “transport”, as well as “chitin catabolic processes”, as to be expected during mycoparasitism (Fig. **6c**). During interaction with the ECM, significantly enriched GO terms were predominantly associated with (extracellular) carbohydrate metabolism and cell wall organization (Fig. **6d**).

Overall, many processes related to mycoparasitism were upregulated already in the early stage of interaction with the two pathogens, while the presence of *L. bicolor* did not lead to any strong induction or repression of specific genes. Also during DC, only contact with the plant pathogens led to clearly recognizable antagonistic gene expression patterns.

### Transcriptomic alterations in *L. bicolor* in confrontation with *T. harzianum*

To identify ECM genes involved in the interaction with *Trichoderma,* we next investigated transcriptomic changes in *L. bicolor* during MC and DC with *T. harzianum*. The LFC values were similarly small as in *T. harzianum* during interaction with *L. bicolor* (Supplementary Fig. **S3**). PCA analysis of the *L. bicolor* transcriptome revealed very similar expression patterns like in the single fungus controls at 3 days (MC) and only clear distinct clustering during DC (six days; Fig. **4d,e**).

During MC, 117 and 329 DEGs were up- and downregulated, respectively, in *L. bicolor* in confrontation with *T. harzianum* (Fig. **5b**). Enrichment analysis of upregulated DEGs in *L. bicolor* after three days revealed enrichment (*p*-adj. < 0.05) of only one GO term, “DNA binding transcription factor activity” (GO:0003700), while no pathways in FunCat or KEGG were significantly enriched (Supplementary Table **S4**). However, several of the upregulated DEGs were annotated with FunCat main categories “Transcription”, “Metabolism”, “Protein with binding function or cofactor requirement”, and “Cellular communication / signal transduction”. This includes one Nrg1-like Zn-finger transcription factor (LACBIDRAFT_296037) being significantly induced in presence of *T. harzianum*, as well as two SNF2 family DNA-dependent ATPases (LACBIDRAFT_301027, LACBIDRAFT_396054). Notably, three upregulated DEGs (LACBIDRAFT_240638, LACBIDRAFT_246709, LACBIDRAFT_248257) were linked to “G-protein coupled receptor signalling, cellular communication / signal transduction mechanism” pathways, indicating the initiation of signaling cascades in *L. bicolor* as potential response to signaling molecules derived by *T. harzianum*. Moreover, several of the 117 upregulated DEGs were assigned to subcategories of the KEGG main pathways “Metabolism”, such as “Biosynthesis of secondary metabolites”, “Steroid biosynthesis”, “alpha-Linolenic acid metabolism”, “Terpenoid backbone biosynthesis”, and “Fatty acid metabolism”, potentially representing an activation of secondary metabolite-based communication.

Interestingly, the most upregulated gene, LACBIDRAFT_304386 (LFC 0.86), was predicted to be a signal peptide-containing protein (likelihood of 0.99). The third most upregulated DEG was an oligopeptide transporter (LACBIDRAFT_302225; LFC 0.71), suggesting increased fluxes of metabolites. A slightly upregulated signal peptide-containing (likelihood 0.94) tripeptidyl-peptidase II (LACBIDRAFT_191088) might indicate a very moderate triggering of some defense mechanisms. Moreover, a plasma membrane fusion protein (LACBIDRAFT_231982), part of the fusion machinery and involved in stabilizing the plasma membrane (“mating projection tip” GO:0043332; “plasma membrane” GO:0005886) was increased by 39%, which could be indicating a reaction of cell wall and membrane remodeling in response to secreted enzymes or effector proteins by *T. harzianum*.

The DC condition led to a stronger response, with 2314 and 1743 up- and downregulated DEGs (Fig. **5b**), respectively. Upregulated DEGs revealed enriched GO terms of “oxidation-reduction process” (GO:0055114), “oxireductase activity” (GO:0016491), “regulation of transcription” (GO:0006355), and “metal ion binding” (GO:0046872) (Fig. **8a**). From KEGG main category “metabolism”, the pathways of “oxidative phosphorylation”, “citrate cycle (TCA cycle)”, “valine, leucine and isoleucine degradation”, “pyruvate metabolism” and “carbon metabolism” were significantly enriched, and FunCat categories such as “cellular transport, transport facilitation and transport routes” (38.4%) “metabolism” (18.4%), “energy” (15.7%), “protein with binding function of cofactor requirement” (12.8%), “cell rescue, defense and virulence” (11.4%), and “interaction with the environment” (1.8%) (Fig. **8b**). Among the transport-related categories, FunCat descriptions of “endoycytosis”, “drug/toxin transport”, and “ABC transporters” (Fig. **8c**) also indicate defense mechanisms. One of those assigned DEGs was a glutathione transferase (LACBIDRAFT_188517; LFC 5), with known function in detoxification, as well as a multidrug resistance-associated (MDR) ABC transporter (LACBIDRAFT_318236, LFC 2), and a pleiotrophic drug resistance (PDR) ABC transporter (LACBIDRAFT_314719, LFC 2.5).

**Fig. 8:**
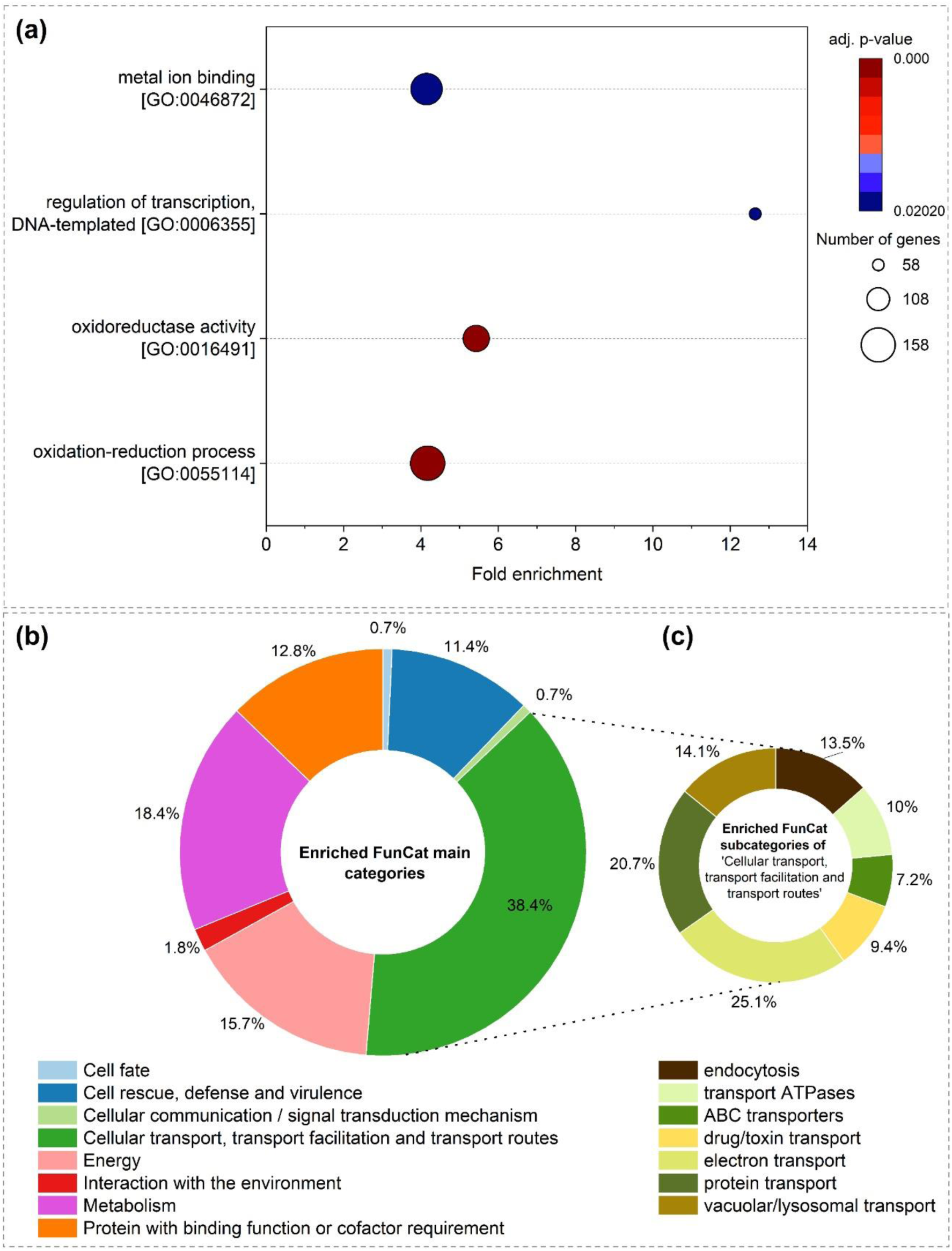
Gene ontology enrichment analysis of upregulated DEGs in *L. bicolor* in presence of *T. harzianum* after six days of co-cultivation (DC) **(a)**. FunCat enrichment analysis of upregulated DEGs in *L. bicolor* during DC (six days) with *T. harzianum*. Significant enriched FunCat main categories **(b)** and enriched subcategories of “Cellular transport, transport facilitation and transport routes” **(c)**. GO terms and FunCat categories were assumed to be significantly enriched with adjusted *p*-value < 0.05.

Furthermore, several Mycorrhiza-induced Small Secreted Proteins (MiSSPs) such as MISSP6.4 (LACBIDRAFT_316998; LFC 5.57), MISSP16.2 (LACBIDRAFT_333197; LFC 1.9), and MISSP22.4 (LACBIDRAFT_303456; LFC 0.48) were significantly upregulated during “overgrowth-phase”. MiSSPs are known to play various roles in plant-fungus interactions, functioning as signaling molecules or effectors.

## 4 Discussion

Fungi engage in diverse interactions with their environment including nearby organisms. The fungi used in this study (*Trichoderma* spp., *Laccaria* spp., *Hebeloma* spp., *Fusarium* spp., and *Alternaria* spp.) can all be found in the soil and rhizosphere and occupy similar environmental niches, while nevertheless having different lifestyles (Hagn et al., 2003). The interactions between two fungal species can manifest as either antagonistic, parasitic, neutral, or synergistic, each yielding distinct outcomes, both for the fungi as well as for potentially involved further interaction partners, such as plants (Venturi & Keel, 2016; Whipps, 2001). One can assume that it would be in the interest of the plants to maximize the number of plant-beneficial microorganisms in their rhizosphere. *Trichoderma* spp. are particularly interesting in this regard, since mycotrophy is thought to be an ancient trait of the entire genus, while many have been shown to work as plant fertilizers (Druzhinina et al., 2011). However, whether the application of *Trichoderma* comes at the expense of overall fungal biodiversity, including plant-beneficial fungi, or is somewhat selective, is controversial so far. A risk assessment when considering *Trichoderma* spp. as biocontrol measure was therefore suggested already early on, requiring to identify the effect on the target (pathogen) population while evaluating negative effects on non-target (native and plant-beneficial) fungal species, which would have detrimental consequences for the host plant (Brimner & Boland, 2003). *Trichoderma viride* and *T. polysporum*, for example, can have a significant impact on ECM, since these species were found to strongly inhibit mycorrhization of black spruce seedlings by *L. bicolor* (Summerbell, 1987). Conversely, the application of the *Trichoderma* bio-inoculant ArborGuard™ on *Pinus radiata* seedlings did not adversely affect the ECM colonization in a nursery system, but did also not promote seedling growth (Minchin et al., 2012). Another study investigated the impact of *T. virens* on pre-mycorrhized *Pinus sylvestris* roots. Intriguingly, while a decreased spore germination of *Trichoderma* was monitored in the rhizosphere, indicating the presence of dampening plant- or ECM-derived processes, the overall plant viability was influenced positively (Werner et al., 2002). The described inhibitory effect of ECM towards *Trichoderma* spp. was observed across various *in vitro* and *in planta* experimental setups (e.g. Guo et al., 2019; Minchin et al., 2012; Summerbell, 1987; Werner et al., 2002). A similar inhibitory effect of the related ECM *Laccaria laccata* towards *Trichoderma virens* in co-culture was already described by Werner et al. (2002). Moreover, Summerbell (1987) reported the absence of typical hyphal structures of *T. viride* towards *L. bicolor* during their interaction, and Zadworny et al. (2007) also described a decreased colony area of *T. virens* and *T. harzianum* during co-culture, as well as altered microtubular cytoskeleton structures in the interaction zone of both fungi, with more pronounced effects in the saprotrophic strains, indicating a stress response.

Likely, the variable outcomes of interaction studies are strongly influenced by the experimental setup, such as space and nutrients, which can unintentionally force the interactions into a specific direction. Our results now demonstrate that in the presence of enough space, *Trichoderma* spp. prefer to grow away from ECMs and do not initiate mycoparasitic programs (either at the physiological or molecular level), that occur in the presence of pathogens long before direct physical contact. These results are in support of a hypothesis arguing in favor of regulatory processes that have evolved to maximize the number of plant-beneficial fungi in the rhizosphere.

### *Trichoderma* are attracted by plant-pathogens and avoid ECM

The dual confrontation assays performed in this study revealed intriguing insights into the response of *Trichoderma* towards fungi of different lifestyles. Co-culture systems are representing a forced competition of two organisms to decide on the fate of the limited resources in confined spaces. Notably, *Trichoderma* strains displayed varying inhibitory effects against *F. graminearum*, *A. alternata*, *H. cylindrosporum,* and *L. bicolor* during different contact conditions. *T. harzianum* exhibited stronger inhibitory effects towards *F. graminearum* and *A. alternata*, especially during MC, suggesting its robust potential to produce diffusible, non-volatile secondary metabolites having a significant effect on the growth of plant-pathogenic fungi (Guo et al., 2019; Küçük & Kivanç; Qualhato et al., 2013; Sood et al., 2020).

Fungal interactions are facilitated through the exchange of signaling molecules, which are released and secreted by one fungal species and subsequently perceived by receptors present in the other participating species (Carreras-Villaseñor et al., 2012; Raut et al., 2014; Zeilinger & Atanasova, 2020). Fungal VOCs are organic chemicals with low molecular weight that originate from metabolic processes within the fungus, evaporate easily at moderate temperature (Macías-Rodríguez et al., 2015) and participate in the communication between fungal species (Guo et al., 2019; Guo et al., 2020; Weikl et al., 2016; Wonglom et al., 2020). Several studies suggest that fungal VOC profiles are similar for fungi with comparable lifestyles (El Jaddaoui et al., 2023; Farh & Jeon, 2020; Guo et al., 2020; Guo et al., 2021; Müller, Faubert et al., 2013; Razo-Belmán et al., 2023) and also *Trichoderma* species have been demonstrated to produce several volatiles (many being sesquiterpenes), as well as receive, and respond to VOCs (Guo et al., 2019; Guo et al., 2020; Huang et al., 2022; Macías-Rodríguez et al., 2020; Nosenko et al., 2023; Rajani et al., 2021; Ruangwong et al., 2021). It is already well-known that VOCs are pivotal in orchestrating inter-species and inter-kingdom signaling within the rhizosphere, a dynamic environment, where various organisms interact (Faure et al., 2009; Werner et al., 2016). They exert influence on growth, defense responses, and behavior of other organisms, with some VOCs exhibiting toxic properties (Hacquard & Schadt, 2015; Sivaprakasam Padmanaban et al., 2022). However, the underlying mechanisms behind these effects and perception mechanisms are poorly understood (Hacquard, 2017; Werner et al., 2016).

In line with our results, earlier *in vitro* co-cultivation scenarios of different *Trichoderma* strains, including *T. harzianum* WM24a1, with the ECM *L. bicolor* already revealed a stronger inhibitory effect of the ECM on the biocontrol strain than the other way around (Guo et al., 2019). The highest VOC emission rate was thereby detected when the two fungi were separated by several cm from each other, indicating VOCs to be an important tool for long-range inter-species communication and recognition. The upregulation of several metabolic KEGG pathways associated to “Biosynthesis of secondary metabolites”, “Steroid biosynthesis”, “alpha-Linolenic acid metabolism”, “Terpenoid backbone biosynthesis”, and “Fatty acid metabolism”, which we now identified in *L. bicolor* in our setup, might reflect the ECM’s response to increased VOC emission rates of *T. harzianum* already at a distance.

The introduced olfactometer “race-tube”-like system now allowed us to observe a strongly accentuated directional growth behavior of *Trichoderma* spp. compared to the conventional plate confrontation assays, demonstrating the positive or negative chemotropism in response to different fungal partners already at a distance and over time. In contrast to plate assays, which lack the capability for directional growth selection, the new system facilitates investigations into interactions spanning longer distances, enabling a choice of growth direction, as it would be the case under most natural conditions. The growth direction is influenced by soluble prey-derived molecules that are involved in fungal chemotropism by acting either as a chemoattractant (positive chemotropism) or as a chemorepellent (negative chemotropism) (Kullnig et al., 2000; Moreno-Ruiz et al., 2021). Our experiments revealed consistent directional growth behavior of both *Trichoderma* strains towards the plant pathogens and away from ECM, even in the absence of soluble biochemicals or metabolites, suggesting a significant role for VOCs in shaping the overall perception and observed physiological response. This underscores the importance of VOCs as potent mediators within the system, exerting a substantial impact on the inter-species interactions. Therefore, the developed olfactometer “race tube”-like system now enables further investigation of VOC-based fungal interactions and perceptions to screen for chemotropic growth.

### Transcriptomic differences in *T. harzianum* and *L. bicolor*

The complex process of mycoparasitism is initiated by prey recognition, directed chemotrophic growth towards the prey, followed by direct attack through (bio-)chemical and physical mechanisms, ultimately leading to death and nutrient release (Chet et al., 1981; Harman, 2006; Moreno-Ruiz et al., 2020; Steindorff et al., 2014). Importantly, the process is signal-dependent, relying on specific inter-species recognition mechanisms (Moreno-Ruiz et al., 2020; Sarma et al., 2014; Zeilinger & Omann, 2007). As the fungi are growing towards each other, they are constantly sensing their environment, and specific downstream signalling cascades are initiated based on the given abiotic and biotic conditions (Bahn et al., 2007; Hinterdobler et al., 2021; Turrà & Di Pietro, 2015; Zeilinger & Atanasova, 2020). Transcriptomic analysis revealed plenty of common DEGs in *T. harzianum* during confrontation with the pathogens, while the confrontation with *L. bicolor* led to distinct - and much more moderate - expression patterns. These observations confirm the lifestyle-specific recognition between the fungi on the molecular level, leading to staging of a rapid and conserved mycoparasitic attack in case of the pathogens and literally a much more “relaxed” communication process in case of the ECM. *Trichoderma* spp. produce extracellular chitinases and proteases in a constitutive manner, leading to enzymatically released chito-oligosaccharides and oligo-peptides in the presence of potential prey, which are sensed and initiate the expression of mycoparasitism-related genes (Benítez et al., 2005; Brunner et al., 2003; Druzhinina et al., 2011; las Mercedes Dana et al., 2001; Seidl et al., 2009). During confrontation with *A. alternata* and *F. graminearum* three chitinases, belonging to the hydrolase family 18, were upregulated in *T. harzianum* already at distance (three days, MC), while the presence of *L. bicolor* did not activate these genes. Additionally, several genes annotated with hydrolase activity, such as an endo-1,3-β-D-glucanase (M431DRAFT_479664) and a peptidase S1 domain-containing protein (M431DRAFT_526221), presumably involved in fungal cell wall degradation and proteolysis, were induced during confrontations with *A. alternata* and *F. graminearum*, but downregulated during confrontation with *L. bicolor*. These results are in line with another comparative transcriptomic analysis that revealed a clear upregulation of mycoparasitism-related genes in *T. atroviride*, *T. reesei*, *T. virens,* as well as *T. harzianum* before direct physical contact with a fungal prey (Atanasova et al., 2013; Wang et al., 2024). The absence of induction of those genes during confrontation with *L. bicolor* is speaking in favour of a non-aggressive interaction. Nonetheless, the question whether this is the result of a self-governed decision by *Trichoderma* or is enforced by repressive or inhibitory metabolites produced by *L. bicolor* remains open so far. However, *L. bicolor* displayed only minimal transcriptomic changes in this situation, suggesting a lack of aggressive adaptations.

Next to VOCs, signalling molecules and prey-derived oligo-peptides are assumed to act as ligands for G-protein-coupled receptors (GPCRs) that are involved in transduction of extracellular signals to intracellular-signaling networks in fungi (Xue et al., 2008) and part of the prey-sensing cascade in *Trichoderma* (Brunner et al., 2008; Hinterdobler et al., 2021; Lin et al., 2019; Seidl et al., 2009; Zeilinger & Atanasova, 2020). Signal reception triggers downstream events via signal transduction mechanisms (Bahn et al., 2007; Carreras-Villaseñor et al., 2012), which play a pivotal role in orchestrating the expression of specific sets of genes that govern the ultimate outcome of the interaction between two fungal species (Sarma et al., 2014). During MC, we identified three upregulated genes (LACBIDRAFT_240638; LACBIDRAFT_246709; LACBIDRAFT_248257) in *L. bicolor* in presence of *T. harzianum*, which are annotated with FunCat categories “GPCR signalling”, “cellular communication”, and “signal transduction mechanism”. Furthermore, the upregulation of transcription factor Nrg1 in *L. bicolor*, has a putative function in carbon catabolite repression (Daguerre et al., 2017) and is involved in gene regulation of stress-response to salt and oxidative stress in *Saccaromyces cerevisiae* and *Ustilago maydis* (Sánchez-Arreguin et al., 2021).

Also in *T. harzianum*, a rhodopsin domain-containing protein (M431DRAFT_155394) belonging to the GPCR rhodopsin family A (Harmar, 2001), was significantly downregulated in presence of *L. bicolor* and induced during confrontation with the pathogens. Additionally, a 3’,5’-cyclic-nucleotide phosphodiesterase (M431DRAFT_74093) was found to be significantly upregulated in *T. harzianum* during MC with *L. bicolor*, indicating modulation of intracellular levels of cyclic nucleotides, such as cyclic adenosine 3’,5’ monophosphate (cAMP). Those messenger molecules are synthesized from ATP by adenylate cyclase activity and activate the cAMP-dependent protein kinase, leading to gene expression regulation by phosphorylation of e.g. transcription factors (Sun et al., 2022). Biogenic VOCs emitted by post-harvested tomatoes, specifically ethylene and benzaldehyde, were identified as active compounds found to be tightly bound to GPCRs in *B. cinerea,* leading to a lack of signal transduction to the cAMP pathway, resulting in reduced pathogenicity of the plant-pathogen (Lin et al., 2019). The detected differential regulation of signal transduction-related genes in *L. bicolor* and *T. harzianum*, as well as the altered VOC emission profiles observed by Guo et al. (2019) during co-cultivation emphasize a VOC-mediated inter-species interaction and GPCRs signal transduction, with a distinct outcome of negative chemotropism and no induction of mycoparasitism-related cascades in *T. harzianum*.

Signal-peptide containing proteins have already been described as effectors in beneficial plant-fungus interactions (Plett et al., 2011), as well as in fungus-fungus interspecies interactions (Feldman et al., 2020). Interestingly, we identified the unique upregulation of a small-secreted protein (SSP) (M431DRAFT_96469) in *T. harzianum* in the presence of *L. bicolor*, while it was downregulated during confrontation with *A. alternata* and *F. graminearum* during MC. This SSP is a homologue (>90% sequence identity) of the cysteine-rich effector Tsp1 in *T. virens,* which was found to be induced in the presence of maize (Lamdan et al., 2015) and banana roots (Muthukathan et al., 2020), indicating an involvement in *Trichoderma*-plant interaction and plant defence modulation. Gupta et al. (2021) analysed the function of Tsp1 in *T. virens* and identified structural similarity with the two fungal effector proteins PevD1 (Bu et al., 2014) and Alt a1 (Chruszcz et al., 2012), which are both interacting with plant defence proteins. The upregulation of this effector in *T. harzianum* confrontation with *L. bicolor* is a new finding and suggests an additional involvement in fungal inter-species interactions.

Also in *L. bicolor,* several short signal peptide-containing proteins were significantly upregulated during confrontation with *T. harzianum*, which might modulate the interaction by acting as secreted effector proteins. While ECM fungi are characterized by a restricted number of carbohydrate-active enzymes (CAZymes), their repertoires of secreted proteases and lipases is similar to saprotrophic fungi, and their secretomes are enriched in SSPs (Pellegrin et al., 2015). Further studies are needed to investigate the role of those detected and potentially secreted proteins in *L. bicolor*. Moreover, considering the plant roots as holobiont, further investigation into the influence of the host plant on the interaction of *T. harzianum* and *L. bicolor* on the mycorrhized roots is required to gain further insights into the fungus-fungus signaling mechanisms and the influence of the plant on the overall outcome of the tripartite interaction.

Concluding, we explored the complex interactions between *Trichoderma* spp. and fungal partners of different lifestyles, including two plant-beneficial ectomycorrhizal fungi (*L. bicolor* and *H. cylindrosporum*), and two plant-pathogenic fungi (*F. graminearum* and *A. alternata*), all potentially interacting within the rhizosphere of poplars. The present work allows insight into the distinct interactions of *Trichoderma* with ECM or pathogens, and sheds light on the multifaceted responses of *Trichoderma* towards root-associated fungi of different lifestyles, speaking in favor of a clear potential to distinguish between plant’s friend and foes during mycoparasitic confrontations. The described phenomenon of ECM avoidance already at a distance highlights the potential of *Trichoderma* spp. as a promising biocontrol agent and plant biofertilizer, while emphasizing the complexities of its interactions with various fungal associates.

## Acknowledgments

The authors sincerely thank Karin Pritsch (Helmholtz Munich, Research Unit Environmental Simulation) for valuable advice and Sascha Schäuble (Department of Microbiome Dynamics, Hans-Knöll-Institute) for support with gene ontology analysis. Furthermore, we thank Franziska Vorwerk for excellent technical assistance.

## Competing interests

The authors declare that they have no competing interests.

## Author contributions

JPB, MR, PS and TK planned and designed the research. PS performed the experiments. PS, PBSP and JK analyzed data. PS wrote the manuscript. JPB, TK, MR and JPS reviewed and edited the manuscript.

## Data availability

Raw sequencing data is deposited on NCBI SRA server and can be accessed under BioProject number PRJNA1100411 and BioSample numbers SAMN40968214-SAMN40968225.

## Funding

The project was supported by Deutsche Forschungsgemeinschaft (DFG project number BE 6069/4-1 to PhB and DFG project number RO 6311/4-1 to MR)

## Notes

### Competing Interest Statement

The authors have declared no competing interest.

## 5 References

Abbey, J. A., Percival, D., Abbey, Asiedu, S. K., Prithiviraj, B., & Schilder, A. (2019). Biofungicides as alternative to synthetic fungicide control of grey mould (*Botrytis cinerea*) – prospects and challenges. Biocontrol Science and Technology, 29(3), 207–228. 10.1080/09583157.2018.1548574

Afzal, I., Sabir, A., & Sikandar, S. (2021). *Trichoderma*: Biodiversity, Abundances, and Biotechnological Applications. In A. N. Yadav (Ed.), Fungal Biology. Recent Trends in Mycological Research (pp. 293–315). Springer International Publishing. 10.1007/978-3-030-60659-6_13

Aleandri, M. P., Chilosi, G., Bruni, N., Tomassini, A., Vettraino, A. M., & Vannini, A. (2015). Use of nursery potting mixes amended with local *Trichoderma* strains with multiple complementary mechanisms to control soil-borne diseases. Crop Protection, 67, 269–278. 10.1016/j.cropro.2014.10.023

Alfiky, A., & Weisskopf, L. (2021). Deciphering *Trichoderma*-Plant-Pathogen Interactions for Better Development of Biocontrol Applications. Journal of Fungi (Basel, Switzerland), 7(1). 10.3390/jof7010061

Almagro Armenteros, J. J., Tsirigos, K. D., Sønderby, C. K., Petersen, T. N., Winther, O., Brunak, S., Heijne, G. von, & Nielsen, H. (2019). Signalp 5.0 improves signal peptide predictions using deep neural networks. Nature Biotechnology, 37(4), 420–423. 10.1038/s41587-019-0036-z

Asef, M., Goltapeh, E., & Danesh, Y. (2008). Antagonistic Effects of *Trichoderma* Species in Biocontrol of *Armillaria Mellea* in Fruit Trees in Iran. Journal of Plant Protection Research, 48(2), 213–222. 10.2478/v10045-008-0025-6

Atanasova, L., Le Crom, S., Gruber, S., Coulpier, F., Seidl-Seiboth, V., Kubicek, C. P., & Druzhinina, I. S. (2013). Comparative transcriptomics reveals different strategies of *Trichoderma* mycoparasitism. BMC Genomics, 14, 121. 10.1186/1471-2164-14-121

Bahn, Y.-S., Xue, C., Idnurm, A., Rutherford, J. C., Heitman, J., & Cardenas, M. E. (2007). Sensing the environment: Lessons from fungi. Nature Reviews. Microbiology, 5(1), 57–69. 10.1038/nrmicro1578

Behnke, K., Ehlting, B., Teuber, M., Bauerfeind, M., Louis, S., Hänsch, R., Polle, A., Bohlmann, J., & Schnitzler, J.-P. (2007). Transgenic, non-isoprene emitting poplars don’t like it hot. The Plant Journal: For Cell and Molecular Biology, 51(3), 485–499. 10.1111/j.1365-313X.2007.03157.x

Benítez, T., Rincón, A., Limón, M. C., & Codón, A. (2005). Biocontrol mechanism of *Trichoderma* strains. International Microbiology: The Official Journal of the Spanish Society for Microbiology, 7, 249–260.

Bonfante, P., & Genre, A. (2010). Mechanisms underlying beneficial plant-fungus interactions in mycorrhizal symbiosis. Nature Communications, 1, 48. 10.1038/ncomms1046

Brimner, T. A., & Boland, G. J. (2003). A review of the non-target effects of fungi used to biologically control plant diseases. Agriculture, Ecosystems & Environment, 100(1), 3–16. 10.1016/S0167-8809(03)00200-7

Brunner, K., Omann, M., Pucher, M. E., Delic, M., Lehner, S. M., Domnanich, P., Kratochwill, K., Druzhinina, I., Denk, D., & Zeilinger, S. (2008). *Trichoderma* G protein-coupled receptors: Functional characterisation of a cAMP receptor-like protein from *Trichoderma atroviride*. Current Genetics, 54(6), 283–299. 10.1007/s00294-008-0217-7

Brunner, K., Peterbauer, C. K., Mach, R. L., Lorito, M., Zeilinger, S., & Kubicek, C. P. (2003). The Nag1 N-acetylglucosaminidase of *Trichoderma atroviride* is essential for chitinase induction by chitin and of major relevance to biocontrol. Current Genetics, 43(4), 289–295. 10.1007/s00294-003-0399-y

Bu, B., Qiu, D., Zeng, H., Guo, L., Yuan, J., & Yang, X. (2014). A fungal protein elicitor PevD1 induces *Verticillium* wilt resistance in cotton. Plant Cell Reports, 33(3), 461–470. 10.1007/s00299-013-1546-7

Cai, F., & Druzhinina, I. S. (2021). In honor of John Bissett: authoritative guidelines on molecular identification of *Trichoderma*. Fungal Diversity, 107(1), 1–69. 10.1007/s13225-020-00464-4

Carreras-Villaseñor, N., Sánchez-Arreguín, J. A., & Herrera-Estrella, A. (2012). *Trichoderma*: Sensing the environment for survival and dispersal. Microbiology, 158(Pt 1), 3–16. 10.1099/mic.0.052688-0

Chet, I., Harman, G. E., & Baker, R. (1981). *Trichoderma hamatum*: Its hyphal interactions with *Rhizoctonia solani* and *Pythium* spp. Microbial Ecology, 7(1), 29–38. 10.1007/BF02010476

Chet, I., Ralph R. Baker, & Peter E. Dunn (1990). Mycoparasitism - recognition, physiology and ecology. In

Chruszcz, M., Chapman, M. D., Osinski, T., Solberg, R., Demas, M., Porebski, P. J., Majorek, K. A., Pomés, A., & Minor, W. (2012). Alternaria alternata allergen Alt a 1: A unique β-barrel protein dimer found exclusively in fungi. The Journal of Allergy and Clinical Immunology, 130(1), 241–7.e9. 10.1016/j.jaci.2012.03.047

Daguerre, Y., Levati, E., Ruytinx, J., Tisserant, E., Morin, E., Kohler, A., Montanini, B., Ottonello, S., Brun, A., Veneault-Fourrey, C., & Martin, F. (2017). Regulatory networks underlying mycorrhizal development delineated by genome-wide expression profiling and functional analysis of the transcription factor repertoire of the plant symbiotic fungus *Laccaria bicolor*. BMC Genomics, 18(1), 737. 10.1186/s12864-017-4114-7

Di Tommaso, P., Chatzou, M., Floden, E. W., Barja, P. P., Palumbo, E., & Notredame, C. (2017). Nextflow enables reproducible computational workflows. Nature Biotechnology, 35(4), 316–319. 10.1038/nbt.3820

Dobin, A., Davis, C. A., Schlesinger, F., Drenkow, J., Zaleski, C., Jha, S., Batut, P., Chaisson, M., & Gingeras, T. R. (2013). Star: Ultrafast universal RNA-seq aligner. Bioinformatics (Oxford, England), 29(1), 15–21. 10.1093/bioinformatics/bts635

Druzhinina, I. S., Seidl-Seiboth, V., Herrera-Estrella, A., Horwitz, B. A., Kenerley, C. M., Monte, E., Mukherjee, P. K., Zeilinger, S., Grigoriev, I., & Kubicek, C. P. (2011). *Trichoderma*: The genomics of opportunistic success. Nature Reviews. Microbiology, 9(10), 749–759. 10.1038/nrmicro2637

El Jaddaoui, I., Rangel, D. E. N., & Bennett, J. W. (2023). Fungal volatiles have physiological properties. Fungal Biology, 127(7-8), 1231–1240. 10.1016/j.funbio.2023.03.005

Ewels, P. A., Peltzer, A., Fillinger, S., Patel, H., Alneberg, J., Wilm, A., Garcia, M. U., Di Tommaso, P., & Nahnsen, S. (2020). The nf-core framework for community-curated bioinformatics pipelines. Nature Biotechnology, 38(3), 276–278. 10.1038/s41587-020-0439-x

Farh, M. E.-A., & Jeon, J. (2020). Roles of Fungal Volatiles from Perspective of Distinct Lifestyles in Filamentous Fungi. The Plant Pathology Journal, 36(3), 193–203. 10.5423/PPJ.RW.02.2020.0025

Faure, D., Vereecke, D., & Leveau, J. H. J. (2009). Molecular communication in the rhizosphere. Plant and Soil, 321(1-2), 279–303. 10.1007/s11104-008-9839-2

Feldman, D., Yarden, O., & Hadar, Y. (2020). Seeking the Roles for Fungal Small-Secreted Proteins in Affecting Saprophytic Lifestyles. Frontiers in Microbiology, 11, 455. 10.3389/fmicb.2020.00455

Frąc, M., Hannula, S. E., Bełka, M., & Jędryczka, M. (2018). Fungal Biodiversity and Their Role in Soil Health. Frontiers in Microbiology, 9, 707. 10.3389/fmicb.2018.00707

Garnica-Vergara, A., Barrera-Ortiz, S., Muñoz-Parra, E., Raya-González, J., Méndez-Bravo, A., Macías-Rodríguez, L., Ruiz-Herrera, L. F., & López-Bucio, J. (2016). The volatile 6-pentyl-2H-pyran-2-one from *Trichoderma atroviride* regulates *Arabidopsis thaliana* root morphogenesis via auxin signaling and ETHYLENE INSENSITIVE 2 functioning. The New Phytologist, 209(4), 1496–1512. 10.1111/nph.13725

Grodnitskaya, I. D., & Sorokin, N. D. (2006). Use of micromycetes *Trichoderma* for soil bioremediation in tree nurseries. Biology Bulletin, 33(4), 400–403. 10.1134/S1062359006040121

Guo, Y., Ghirado, A., Weber, B., Schnitzler, J.-P., Benz, J. P., & Rosenkranz, M. (2019). *Trichoderma* Species Differ in Their Volatile Profiles and in Antagonism Towards Ectomycorrhiza *Laccaria bicolor*. Frontiers in Microbiology, 10, 891. 10.3389/fmicb.2019.00891

Guo, Y., Jud, W., Ghirardo, A., Antritter, F., Benz, J. P., Schnitzler, J.-P., & Rosenkranz, M. (2020). Sniffing fungi - phenotyping of volatile chemical diversity in *Trichoderma* species. The New Phytologist, 227(1), 244–259. 10.1111/nph.16530

Guo, Y., Jud, W., Weikl, F., Ghirardo, A., Junker, R. R., Polle, A., Benz, J. P., Pritsch, K., Schnitzler, J.-P., & Rosenkranz, M. (2021). Volatile organic compound patterns predict fungal trophic mode and lifestyle. Communications Biology, 4(1), 673. 10.1038/s42003-021-02198-8

Gupta, G. D., Bansal, R., Mistry, H., Pandey, B., & Mukherjee, P. K. (2021). Structure-function analysis reveals *Trichoderma virens* Tsp1 to be a novel fungal effector protein modulating plant defence. International Journal of Biological Macromolecules, 191, 267–276. 10.1016/j.ijbiomac.2021.09.085

Guzmán-Guzmán, P., Porras-Troncoso, M. D., Olmedo-Monfil, V., & Herrera-Estrella, A. (2019). *Trichoderma* Species: Versatile Plant Symbionts. Phytopathology, 109(1), 6–16. 10.1094/PHYTO-07-18-0218-RVW

Hacquard, S. (2017). Commentary: Microbial Small Talk: Volatiles in Fungal-Bacterial Interactions. Frontiers in Microbiology, 8, 1. 10.3389/fmicb.2017.00001

Hacquard, S., Petre, B., Frey, P., Hecker, A., Rouhier, N., & Duplessis, S. (2011). The poplar-poplar rust interaction: Insights from genomics and transcriptomics. Journal of Pathogens, 2011, 716041. 10.4061/2011/716041

Hacquard, S., & Schadt, C. W. (2015). Towards a holistic understanding of the beneficial interactions across the *Populus* microbiome. The New Phytologist, 205(4), 1424– 1430. 10.1111/nph.13133

Hagn, A., Pritsch, K., Schloter, M., & Munch, J. C. (2003). Fungal diversity in agricultural soil under different farming management systems, with special reference to biocontrol strains of *Trichoderma* spp. Biology and Fertility of Soils, 38(4), 236–244. 10.1007/s00374-003-0651-0

Harman, G. E. (2006). Overview of Mechanisms and Uses of *Trichoderma* spp. Phytopathology, 96(2), 190–194. 10.1094/PHYTO-96-0190

Harman, G. E., Howell, C. R., Viterbo, A., Chet, I., & Lorito, M. (2004). *Trichoderma* species-opportunistic, avirulent plant symbionts. Nature Reviews. Microbiology, 2(1), 43–56. 10.1038/nrmicro797

Harmar, A. J. (2001). Family-B G-protein-coupled receptors. Genome Biology, 2(12), REVIEWS3013. 10.1186/gb-2001-2-12-reviews3013

Harris, M. A., Clark, J., Ireland, A., Lomax, J., Ashburner, M., Foulger, R., Eilbeck, K., Lewis, S., Marshall, B., Mungall, C., Richter, J., Rubin, G. M., Blake, J. A., Bult, C., Dolan, M., Drabkin, H., Eppig, J. T., Hill, D. P., Ni, L., … White, R. (2004). The Gene Ontology (GO) database and informatics resource. Nucleic Acids Research, 32(Database issue), D258–61. 10.1093/nar/gkh036

Hassani, M. A., Durán, P., & Hacquard, S. (2018). Microbial interactions within the plant holobiont. Microbiome, 6(1), 58. 10.1186/s40168-018-0445-0

Hermosa, R., Rubio, M. B., Cardoza, R. E., Nicolás, C., Monte, E., & Gutiérrez, S. (2013). The contribution of *Trichoderma* to balancing the costs of plant growth and defense. International Microbiology, 16(2), 69–80. 10.2436/20.1501.01.181

Hewitt, E. J., & Smith, T. A. (1975). Plant Mineral Nutrition. Journal of Plant Nutrition and Soil Science, 142(6), 875. 10.1002/jpln.19791420613

Hinterdobler, W., Li, G., Turrà, D., Schalamun, M., Kindel, S., Sauer, U., Beier, S., Iglesias, A. R., Compant, S., Vitale, S., Di Pietro, A., & Schmoll, M. (2021). Integration of chemosensing and carbon catabolite repression impacts fungal enzyme regulation and plant associations. BioRxiv. Advance online publication. 10.1101/2021.05.06.442915

Howe, E. A., Sinha, R., Schlauch, D., & Quackenbush, J. (2011). Rna-Seq analysis in MeV. Bioinformatics (Oxford, England), 27(22), 3209–3210. 10.1093/bioinformatics/btr490

Huang, R., Zhou, R., Zhou, S., Lin, H., Lu, S., Qiu, J., & He, J. (2022). New sesquiterpene from a soil fungus of Trichoderma sp. Natural Product Research, 1–8. 10.1080/14786419.2022.2159398

Hyakumachi, M., & Kubota, M. (2004). Fungi as plant growth promoter and disease suppressor. Fungal Biotechnology in Agricultural, Food, and Environmental Applications, 21, 101–110.

Kanehisa, M., & Goto, S. (2000). Kegg: Kyoto encyclopedia of genes and genomes. Nucleic Acids Research, 28(1), 27–30. 10.1093/nar/28.1.27

Karliński, L., Rudawska, M., Kieliszewska-Rokicka, B., & Leski, T. (2010). Relationship between genotype and soil environment during colonization of poplar roots by mycorrhizal and endophytic fungi. Mycorrhiza, 20(5), 315–324. 10.1007/s00572-009-0284-8

Küçük, Ç., & Kivanç, M. Isolation of *Trichoderma* Spp. and Determination of Their Antifungal, Biochemical and Physiological Features.

Kullnig, C., Mach, R. L., Lorito, M., & Kubicek, C. P. (2000). Enzyme diffusion from *Trichoderma atroviride* (= *T. Harzianum* P1) to *Rhizoctonia solani* is a prerequisite for triggering of *Trichoderma* ech42 gene expression before mycoparasitic contact. Applied and Environmental Microbiology, 66(5), 2232–2234. 10.1128/AEM.66.5.2232-2234.2000

Kwaśna, H., Szewczyk, W., Baranowska, M., Gallas, E., Wiśniewska, M., & Behnke-Borowczyk, J. (2021). Mycobiota Associated with the Vascular Wilt of Poplar. Plants (Basel, Switzerland), 10(5). 10.3390/plants10050892

Lamdan, N.-L., Shalaby, S., Ziv, T., Kenerley, C. M., & Horwitz, B. A. (2015). Secretome of *Trichoderma* interacting with maize roots: Role in induced systemic resistance. Molecular & Cellular Proteomics : MCP, 14(4), 1054–1063. 10.1074/mcp.M114.046607

las Mercedes Dana, M. de, Limón, M. C., Mejías, R., Mach, R. L., Benítez, T., Pintor-Toro, J. A., & Kubicek, C. P. (2001). Regulation of chitinase 33 (*chit33*) gene expression in *Trichoderma harzianum*. Current Genetics, 38(6), 335–342. 10.1007/s002940000169

Lin, Y [Yongwen], Ruan, H., Akutse, K. S., Lai, B., Lin, Y [Yizhang], Hou, Y., & Zhong, F. (2019). Ethylene and Benzaldehyde Emitted from Postharvest Tomatoes Inhibit Botrytis cinerea via Binding to G-Protein Coupled Receptors and Transmitting with cAMP-Signal Pathway of the Fungus. Journal of Agricultural and Food Chemistry, 67(49), 13706–13717. 10.1021/acs.jafc.9b05778

Liu, Y., He, P [Pengbo], He, P [Pengfei], Munir, S., Ahmed, A., Wu, Y., Yang, Y., Lu, J., Wang, J [Jiansong], Yang, J., Pan, X., Tian, Y., & He, Y. (2022). Potential biocontrol efficiency of *Trichoderma* species against oomycete pathogens. Frontiers in Microbiology, 13, 974024. 10.3389/fmicb.2022.974024

López-Bucio, J., Ramón Pelagio-Flores, & Alfredo Herrera-Estrella (2015). *Trichoderma* as biostimulant: exploiting the multilevel properties of a plant beneficial fungus. Scientia Horticulturae, 196, 109–123. 10.1016/j.scienta.2015.08.043

Love, M. I., Huber, W., & Anders, S. (2014). Moderated estimation of fold change and dispersion for RNA-seq data with DESeq2. Genome Biology, 15(12), 550. 10.1186/s13059-014-0550-8

Luo, Z.-B., Janz, D., Jiang, X., Göbel, C., Wildhagen, H., Tan, Y., Rennenberg, H., Feussner, I., & Polle, A. (2009). Upgrading root physiology for stress tolerance by ectomycorrhizas: Insights from metabolite and transcriptional profiling into reprogramming for stress anticipation. Plant Physiology, 151(4), 1902–1917. 10.1104/pp.109.143735

Ma, L., Wu, X., & Zheng, L. (2008). Mycorrhizal formation of nine ectomycorrhizal fungi on poplar cuttings. Frontiers of Forestry in China, 3(4), 475–479. 10.1007/s11461-008-0077-9

Macías-Rodríguez, L., Contreras-Cornejo, H. A., Adame-Garnica, S. G., Del-Val, E., & Larsen, J. (2020). The interactions of *Trichoderma* at multiple trophic levels: Inter-kingdom communication. Microbiological Research, 240, 126552. 10.1016/j.micres.2020.126552

Macías-Rodríguez, L., Contreras-Cornejo, H. Á., López-Bucio, J. S., & López-Bucio, J. (2015). Recent advancements in the role of volatile organic compounds from fungi. In V. K. Gupta, R. L. Mach, & S. Sreenivasaprasad (Eds.), Fungal Biomolecules (pp. 87–99). Wiley. 10.1002/9781118958308.ch7

Manzar, N., Kashyap, A. S., Goutam, R. S., Rajawat, M. V. S., Sharma, P. K., Sharma, S. K., & Singh, H. V. (2022). *Trichoderma*: Advent of Versatile Biocontrol Agent, Its Secrets and Insights into Mechanism of Biocontrol Potential. Sustainability, 14(19), 12786. 10.3390/su141912786

Martin, M. (2011). Cutadapt removes adapter sequences from high-throughput sequencing reads. EMBnet.Journal, 17(1), 10. 10.14806/ej.17.1.200

Mayor, J., Bahram, M., Henkel, T., Buegger, F., Pritsch, K., & Tedersoo, L. (2015). Ectomycorrhizal impacts on plant nitrogen nutrition: Emerging isotopic patterns, latitudinal variation and hidden mechanisms. Ecology Letters, 18(1), 96–107. 10.1111/ele.12377

Minchin, R. F., Ridgway, H. J., Condron, L., & Jones, E. E. (2012). Influence of inoculation with a *Trichoderma* bio-inoculant on ectomycorrhizal colonisation of *Pinus radiata* seedlings. Annals of Applied Biology, 161(1), 57–67. 10.1111/j.1744-7348.2012.00552.x

Moreno-Ruiz, D., Lichius, A., Turrà, D., Di Pietro, A., & Zeilinger, S. (2020). Chemotropism Assays for Plant Symbiosis and Mycoparasitism Related Compound Screening in *Trichoderma atroviride*. Frontiers in Microbiology, 11, 601251. 10.3389/fmicb.2020.601251

Moreno-Ruiz, D., Salzmann, L., Fricker, M. D., Zeilinger, S., & Lichius, A. (2021). Stress-Activated Protein Kinase Signalling Regulates Mycoparasitic Hyphal-Hyphal Interactions in *Trichoderma atroviride*. Journal of Fungi (Basel, Switzerland), 7(5). 10.3390/jof7050365

Müller, A., Faubert, P., Hagen, M., Castell, W. zu, Polle, A., Schnitzler, J.-P., & Rosenkranz, M. (2013). Volatile profiles of fungi-chemotyping of species and ecological functions. Fungal Genetics and Biology : FG & B, 54, 25–33. 10.1016/j.fgb.2013.02.005

Müller, A., Volmer, K., Mishra-Knyrim, M., & Polle, A. (2013). Growing poplars for research with and without mycorrhizas. Frontiers in Plant Science, 4, 332. 10.3389/fpls.2013.00332

Muthukathan, G., Mukherjee, P., Salaskar, D., Pachauri, S., Tak, H., Ganapathi, T. R., & Mukherjee, P. K. (2020). Secretome of *Trichoderma virens* induced by banana roots - identification of novel fungal proteins for enhancing plant defence. Physiological and Molecular Plant Pathology, 110, 101476. 10.1016/j.pmpp.2020.101476

Nadziakiewicz, M., Kurzawińska, H., Mazur, S., & Tekielska, D. (2018). *Alternaria alternata* – the main causal agent of disease symptoms in juniper, rose, yew and highbush blueberry in nurseries in southern Poland. Folia Horticulturae, 30(1), 15–25. 10.2478/fhort-2018-0002

Nawrocka, J., & Małolepsza, U. (2013). Diversity in plant systemic resistance induced by *Trichoderma*. Biological Control, 67(2), 149–156. 10.1016/j.biocontrol.2013.07.005

Nosenko, T., Zimmer, I., Ghirardo, A., Köllner, T. G., Weber, B., Polle, A., Rosenkranz, M., & Schnitzler, J.-P. (2023). Predicting functions of putative fungal sesquiterpene synthase genes based on multiomics data analysis. Fungal Genetics and Biology : FG & B, 165, 103779. 10.1016/j.fgb.2023.103779

Nur, A. Z., & Noor, A. B. (2020). Biological functions of *Trichoderma* spp. for agriculture applications. Annals of Agricultural Sciences, 65(2), 168–178. 10.1016/j.aoas.2020.09.003

Oskiera, M., Szczech, M., Stępowska, A., Smolińska, U., & Bartoszewski, G. (2017). Monitoring of *Trichoderma* species in agricultural soil in response to application of biopreparations. Biological Control, 113, 65–72. 10.1016/j.biocontrol.2017.07.005

Ostry, M., Ramstedt, M., Newcombe, G., & Steenackers, M. (2014). Diseases of Poplars and Willows. In J.G. Isebrands and J. Richardson (Ed.), Poplars and Willows: Trees for Society and the Environment (249–76). FAO UN and CABI.

Patro, R., Duggal, G., Love, M. I., Irizarry, R. A., & Kingsford, C. (2017). Salmon provides fast and bias-aware quantification of transcript expression. Nature Methods, 14(4), 417–419. 10.1038/nmeth.4197

Pellegrin, C., Morin, E., Martin, F., & Veneault-Fourrey, C. (2015). Comparative Analysis of Secretomes from Ectomycorrhizal Fungi with an Emphasis on Small-Secreted Proteins. Frontiers in Microbiology, 6, 1278. 10.3389/fmicb.2015.01278

Plett, J. M., Kemppainen, M., Kale, S. D., Kohler, A., Legué, V., Brun, A., Tyler, B. M., Pardo, A. G., & Martin, F. (2011). A secreted effector protein of Laccaria bicolor is required for symbiosis development. Current Biology : CB, 21(14), 1197–1203. 10.1016/j.cub.2011.05.033

Plett, J. M., & Martin, F. (2012). Poplar root exudates contain compounds that induce the expression of MiSSP7 in *Laccaria bicolor*. Plant Signaling & Behavior, 7(1), 12–15. 10.4161/psb.7.1.18357

Polle, A., & Douglas, C. (2010). The molecular physiology of poplars: Paving the way for knowledge-based biomass production. Plant Biology, 12(2), 239–241. 10.1111/j.1438-8677.2009.00318.x

Priebe, S., Kreisel, C., Horn, F., Guthke, R., & Linde, J. (2015). Fungifun2: A comprehensive online resource for systematic analysis of gene lists from fungal species. Bioinformatics (Oxford, England), 31(3), 445–446. 10.1093/bioinformatics/btu627

Przybysz, K., & Przybysz, P. (2013). Poplar wood as raw material for the paper industry in the twenty-first century. Annals of Warsaw University of Life Sciences-SGGW. Forestry and Wood Technology(84).

Qualhato, T. F., Lopes, F. A. C., Steindorff, A. S., Brandão, R. S., Jesuino, R. S. A., & Ulhoa, C. J. (2013). Mycoparasitism studies of *Trichoderma* species against three phytopathogenic fungi: Evaluation of antagonism and hydrolytic enzyme production. Biotechnology Letters, 35(9), 1461–1468. 10.1007/s10529-013-1225-3

Rajani, P., Rajasekaran, C., Vasanthakumari, M. M., Olsson, S. B., Ravikanth, G., & Uma Shaanker, R. (2021). Inhibition of plant pathogenic fungi by endophytic *Trichoderma* spp. Through mycoparasitism and volatile organic compounds. Microbiological Research, 242, 126595. 10.1016/j.micres.2020.126595

Raut, I., Badea-Doni, M., Calin, M., Oancea, F., Vasilescu, G., Sesan, T. E., & Jecu, L. (2014). Effect of volatile and non-volatile metabolites from *Trichoderma* spp. against important phytopathogens. Revista De Chimie, 65(11), 1285–1288.

Razo-Belmán, R., Ángeles-López, Y. I., García-Ortega, L. F., León-Ramírez, C. G., Ortiz-Castellanos, L., Yu, H., & Martínez-Soto, D. (2023). Fungal volatile organic compounds: Mechanisms involved in their sensing and dynamic communication with plants. Frontiers in Plant Science, 14, 1257098. 10.3389/fpls.2023.1257098

Ruangwong, O.-U., Wonglom, P., Suwannarach, N., Kumla, J., Thaochan, N., Chomnunti, P., Pitija, K., & Sunpapao, A. (2021). Volatile Organic Compound from *Trichoderma asperelloides* TSU1: Impact on Plant Pathogenic Fungi. Journal of Fungi (Basel, Switzerland), 7(3). 10.3390/jof7030187

Rubin, E. M. (2008). Genomics of cellulosic biofuels. Nature, 454(7206), 841–845. 10.1038/nature07190

Ruepp, A., Zollner, A., Maier, D., Albermann, K., Hani, J., Mokrejs, M., Tetko, I., Güldener, U., Mannhaupt, G., Münsterkötter, M., & Mewes, H. W. (2004). The FunCat, a functional annotation scheme for systematic classification of proteins from whole genomes. Nucleic Acids Research, 32(18), 5539–5545. 10.1093/nar/gkh894

Rush, T. A., Shrestha, H. K., Gopalakrishnan Meena, M., Spangler, M. K., Ellis., J. C., Labbé, J. L., & Abraham, P. E. (2021). Bioprospecting *Trichoderma*: A Systematic Roadmap to Screen Genomes and Natural Products for Biocontrol Applications. Frontiers in Fungal Biology, 2, Article 716511, 41. 10.3389/ffunb.2021.716511

Sánchez-Arreguin, J. A., Ruiz-Herrera, J., Mares-Rodriguez, F. d. J., León-Ramírez, C. G., Sánchez-Segura, L., Zapata-Morín, P. A., Coronado-Gallegos, J., & Aréchiga-Carvajal, E. T. (2021). Acid pH Strategy Adaptation through NRG1 in Ustilago maydis. Journal of Fungi (Basel, Switzerland), 7(2). 10.3390/jof7020091

Sarma, B. K., Yadav, S. K., Patel, J. S., & Singh, H. B. (2014). Molecular Mechanisms of Interactions of *Trichoderma* with other Fungal Species. The Open Mycology Journal, 8(1), 140–147. 10.2174/1874437001408010140

Schenk, R. U., & Hildebrandt, A. C. (1972). Medium and techniques for induction and growth of monocotyledonous and dicotyledonous plant cell cultures. Canadian Journal of Botany, 50(1), 199–204. 10.1139/b72-026

Schnitzler, J.-P., Louis, S., Behnke, K., & Loivamäki, M. (2010). Poplar volatiles - biosynthesis, regulation and (eco)physiology of isoprene and stress-induced isoprenoids. Plant Biology, 12(2), 302–316. 10.1111/j.1438-8677.2009.00284.x

Seidl, V., Song, L., Lindquist, E., Gruber, S., Koptchinskiy, A., Zeilinger, S., Schmoll, M., Martínez, P., Sun, J., Grigoriev, I., Herrera-Estrella, A., Baker, S. E., & Kubicek, C. P. (2009). Transcriptomic response of the mycoparasitic fungus *Trichoderma atroviride* to the presence of a fungal prey. BMC Genomics, 10, 567. 10.1186/1471-2164-10-567

Sharma, K., Mishra, A. K., & Misra, R. S. (2009). Morphological, Biochemical and Molecular Characterization of Trichoderma harzianum Isolates for their Efficacy as Biocontrol Agents. Journal of Phytopathology, 157(1), 51–56. 10.1111/j.1439-0434.2008.01451.x

Sharma, P., Kumar, V., Ramesh, R., Saravanan, K., Deep, S., Sharma, M., Mahesh, S., & Dinesh, S. (2011). Biocontrol genes from *Trichoderma* species: a review. African Journal of Biotechnology, 10(86), 19898–19907.

Shoresh, M., Harman, G. E., & Mastouri, F. (2010). Induced systemic resistance and plant responses to fungal biocontrol agents. Annual Review of Phytopathology, 48, 21–43. 10.1146/annurev-phyto-073009-114450

Sivaprakasam Padmanaban, P. B., Rosenkranz, M., Zhu, P., Kaling, M., Schmidt, A., Schmitt-Kopplin, P., Polle, A., & Schnitzler, J.-P. (2022). Mycorrhiza-Tree-Herbivore Interactions: Alterations in Poplar Metabolome and Volatilome. Metabolites, 12(2). 10.3390/metabo12020093

Sood, M., Kapoor, D. K. V., Sheteiwy, M. S., Ramakrishnan, M., Landi, M., Arani, F., & Sharma, A. (2020). *Trichoderma*: The “Secrets” of a Multitalented Biocontrol Agent. Plants, 9(6), 729.

Stange, P., Seidl, S., Karl, T., & Benz, J. P. (2023). Evaluation of Trichoderma isolates as biocontrol measure against Claviceps purpurea. European Journal of Plant Pathology, 167(4), 651–675. 10.1007/s10658-023-02716-w

Steindorff, A. S., Soller Ramada, M. H., Guedes Coelho, A. S., Miller, R. N. G., Pappas, G. J., Ulhoa, C. J., & Noronha, E. F. (2014). Identification of mycoparasitism-related genes against the phytopathogen *Sclerotinia sclerotiorum* through transcriptome and expression profile analysis in *Trichoderma harzianum*. BMC Genomics, 15, 204.

Stenberg, J. A., Sundh, I., Becher, P. G., Björkman, C., Dubey, M., Egan, P. A., Friberg, H., Gil, J. F., Jensen, D. F., Jonsson, M., Karlsson, M., Khalil, S., Ninkovic, V., Rehermann, G., Vetukuri, R. R., & Viketoft, M. (2021). When is it biological control? A framework of definitions, mechanisms, and classifications. Journal of Pest Science, 94(3), 665–676. 10.1007/s10340-021-01354-7

Summerbell, R. C. (1987). The inhibitory effect of *Trichoderma* species and other soil microfungi on formation of mycorrhiza by *Laccaria bicolor in vitro*. The New Phytologist, 105(3), 437–448. 10.1111/j.1469-8137.1987.tb00881.x

Sun, Z.-B., Yu, S.-F., Wang, C.-L., & Wang, L. (2022). Camp Signalling Pathway in Biocontrol Fungi. Current Issues in Molecular Biology, 44(6), 2622–2634. 10.3390/cimb44060179

Thambugala, K. M., Daranagama, D. A., Phillips, A. J. L., Kannangara, S. D., & Promputtha, I. (2020). Fungi vs. Fungi in Biocontrol: An Overview of Fungal Antagonists Applied Against Fungal Plant Pathogens. Frontiers in Cellular and Infection Microbiology, 10, 604923. 10.3389/fcimb.2020.604923

Turrà, D., & Di Pietro, A. (2015). Chemotropic sensing in fungus-plant interactions. Current Opinion in Plant Biology, 26, 135–140. 10.1016/j.pbi.2015.07.004

Tyśkiewicz, R., Nowak, A., Ozimek, E., & Jaroszuk-Ściseł, J. (2022). *Trichoderma*: The Current Status of Its Application in Agriculture for the Biocontrol of Fungal Phytopathogens and Stimulation of Plant Growth. International Journal of Molecular Sciences, 23(4), 2329. 10.3390/ijms23042329

Uniyal, K., Chandra, G., Khan, R. U., & Singh, Y. P. (2018). Selection of Potent Isolates from a Population of *Alternaria Alternata*, a Leaf Spot Pathogen of Poplar. American Journal of Applied Mathematics and Statistics, 6(6), 232–238. 10.12691/ajams-6-6-3

Venturi, V., & Keel, C. (2016). Signaling in the Rhizosphere. Trends in Plant Science, 21(3), 187–198. 10.1016/j.tplants.2016.01.005

Viterbo, A., & Horwitz, B. A. (2010). Mycoparasitism. In K. A. Borkovich & D. J. Ebbole (Eds.), Cellular and molecular biology of filamentous fungi (pp. 676–693). ASM Press. 10.1128/9781555816636.ch42

Wang, Y., Wang, J [Jian], Zhu, X., & Wang, W. (2024). Genome and transcriptome sequencing of *Trichoderma harzianum* T4, an important biocontrol fungus of *Rhizoctonia solani*, reveals genes related to mycoparasitism. Canadian Journal of Microbiology, 70(3), 86–101. 10.1139/cjm-2023-0148

Weikl, F., Ghirardo, A., Schnitzler, J.-P., & Pritsch, K. (2016). Sesquiterpene emissions from *Alternaria alternata* and *Fusarium oxysporum*: Effects of age, nutrient availability, and co-cultivation. Scientific Reports, 6, 22152. 10.1038/srep22152.

Werner, A., Zadworny, M., & Idzikowska, K. (2002). Interaction between *Laccaria laccata* and *Trichoderma virens* in co-culture and in the rhizosphere of *Pinus sylvestris* grown in vitro. Mycorrhiza, 12(3), 139–145. 10.1007/s00572-002-0159-8

Werner, S., Polle, A., & Brinkmann, N. (2016). Belowground communication: Impacts of volatile organic compounds (VOCs) from soil fungi on other soil-inhabiting organisms. Applied Microbiology and Biotechnology, 100(20), 8651–8665. 10.1007/s00253-016-7792-1

Whipps, J. M. (2001). Microbial interactions and biocontrol in the rhizosphere. Journal of Experimental Botany, 52(Spec Issue), 487–511. 10.1093/jexbot/52.suppl_1.487

Wonglom, P., Ito, S., & Sunpapao, A. (2020). Volatile organic compounds emitted from endophytic fungus *Trichoderma asperellum* T1 mediate antifungal activity, defense response and promote plant growth in lettuce (*Lactuca sativa*). Fungal Ecology, 43, 100867. 10.1016/j.funeco.2019.100867

Xue, C., Hsueh, Y.-P., & Heitman, J. (2008). Magnificent seven: Roles of G protein-coupled receptors in extracellular sensing in fungi. FEMS Microbiology Reviews, 32(6), 1010– 1032. 10.1111/j.1574-6976.2008.00131.x

Yu, Z., Liu, Z., Zhang, Y., & Wang, Z. (2022). The disease resistance potential of *Trichoderma asperellum* T-Pa2 isolated from Phellodendron amurense rhizosphere soil. Journal of Forestry Research, 33(1), 321–331. 10.1007/s11676-021-01332-w

Zadworny, M., Tuszyńska, S., Samardakiewicz, S., & Werner, A. (2007). Effects of mutual interaction of Laccaria laccata with *Trichoderma harzianum* and *T. Virens* on the morphology of microtubules and mitochondria. Protoplasma, 232(1-2), 45–53. 10.1007/s00709-007-0276-5

Zeilinger, S., & Atanasova, L. (2020). Sensing and regulation of mycoparasitism-relevant processes in *Trichoderma*. In New and Future Developments in Microbial Biotechnology and Bioengineering (pp. 39–55). Elsevier. 10.1016/B978-0-12-819453-9.00002-7

Zeilinger, S., & Omann, M. (2007). *Trichoderma* Biocontrol: Signal Transduction Pathways Involved in Host Sensing and Mycoparasitism. Gene Regulation and Systems Biology, 1, GRSB.S397. 10.4137/GRSB.S397

Zilber-Rosenberg, I., & Rosenberg, E. (2008). Role of microorganisms in the evolution of animals and plants: The hologenome theory of evolution. FEMS Microbiology Reviews, 32(5), 723–735. 10.1111/j.1574-6976.2008.00123.x

