## Supplementary Information for "Lifestyle-specific responses of *Trichoderma* spp. in mycoparasitic confrontations and implications for biocontrol of *Populus* x *canescens*"

Article acceptance date: Click here to enter a date.

The following **Supporting Information** is available for this article:

**Fig. S1** Representative pictures of *T. harzianum* plate confrontations.

**Fig. S2** Experimental set-up of olfactometer "race tube"- like system and test conditions.

**Fig. S3** Volcano plots of differential gene expression analysis.

**Fig. S4** Expression graphs and gene names based on SOTA cluster organization of the 57 common DEGs in *T. harzianum*.

**Table S1** Genes identified as differentially expressed in *T. harzianum* and *L. bicolor* compared to the control conditions.

**Table S2** Common DEGs in *T. harzianum* during MC with *L. bicolor*, *A. alternata* and *F. graminearum* clustered using SOTA based on expression values.

**Table S3** Venny groups of common up- and downregulated DEGs in *T. harzianum*.

**Table S4** Gene ontology enrichment analysis results.

**Methods S1** Cultivation of *P. x canescens.*

**Methods S2** Preparation and setup of olfactometer "race tube"-like system.

**Methods S3** RNA-extraction from fungal confrontations for transcriptomic analysis.

**Fig. S1** Example pictures of the co-cultivation of *T. harzianum* during media contact (MC) and air contact (AC) with *L. bicolor* and *A. alternata* three days after inoculation of *T. harzianum*.

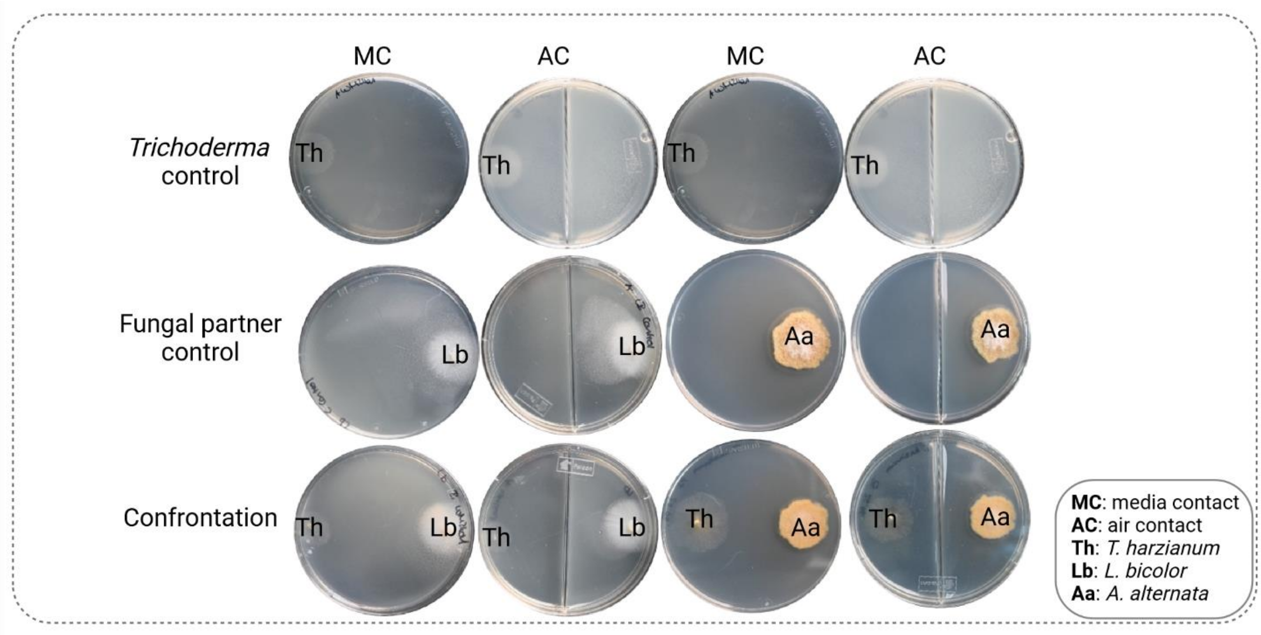

**Fig. S2** Experimental set-up of olfactometer “race tube”- like system and conditions. Two sample cups and one serological pipette were assembled by connecting the two cups with the serological pipette and sealing the joints with physiologically harmless silicon **(a)**. The system is filled with agar-based media in a way that the media level was the same in both tubes and the horizontal serological pipette was filled in half, allowing media and air contact between both test tubes and a flat media surface. Different confrontation scenarios included media contact (MC) and air contact (AC) with *L. bicolor*, *H. cylindrosporum*, *A. alternata*, and *F. graminearum*. For AC, the second fungus was inoculated onto a small petri dish (ø = 6 cm) which was then placed into the second sample cup. As a control, Trichoderma was challenged with itself. Picture of assembled olfactometer “race tube”-like system filled with MMN solid media **(b)**.

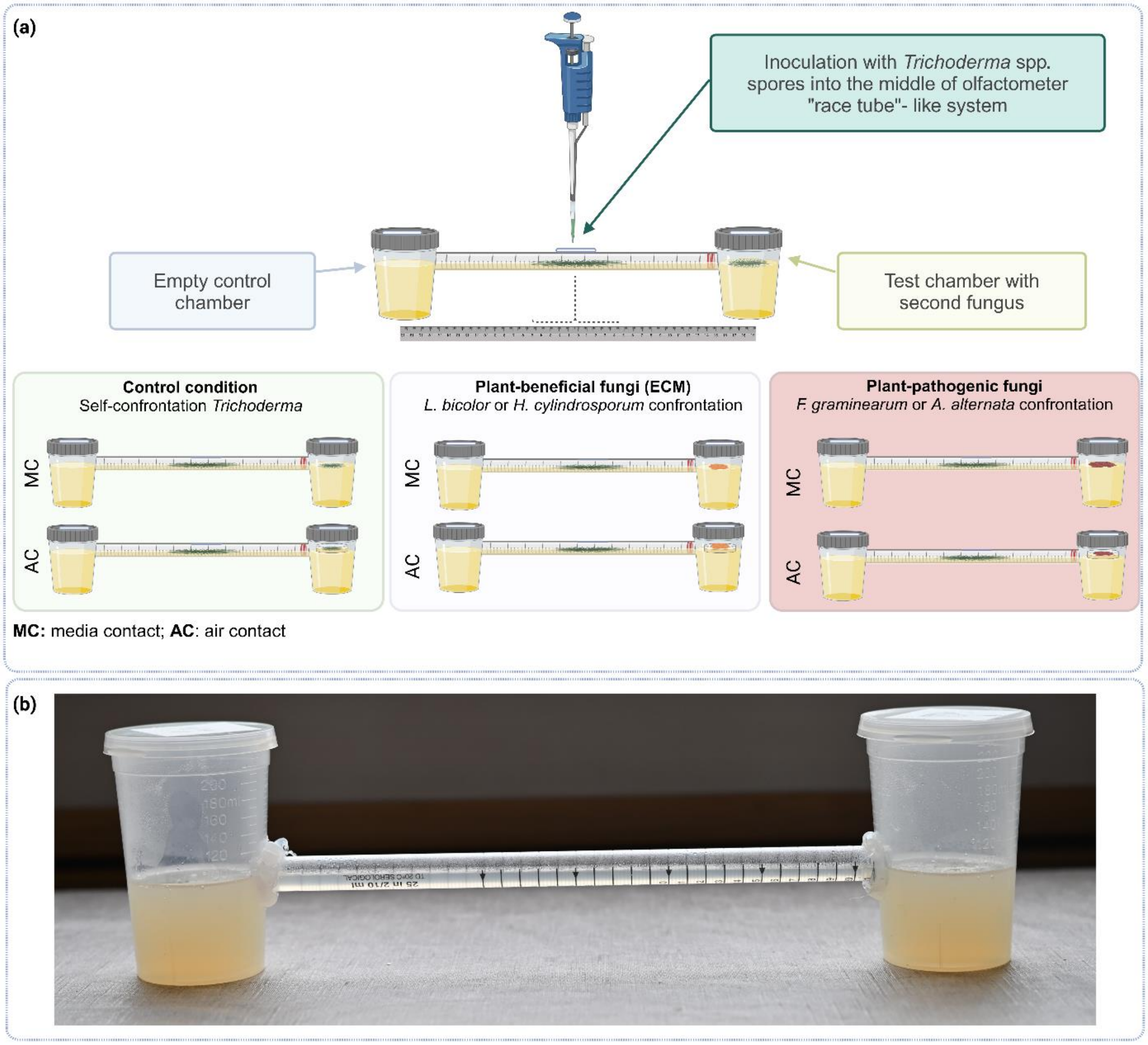

**Fig. S3** Volcano plots of transcriptomic analysis showing all DEGs (orange FDR < 0.05, red FDR < 0.01) with respective FDR (adj. *p*-value) vs. log2 fold change for test conditions vs. control conditions. *T. harzianum* transcriptome during MC interaction with *L. bicolor* TL3 **(a)**, *F. graminearum* TF3 **(b)**, and *A. alternata* TA3 **(c)** after three days normalized to *T. harzianum* control (TC3) and during DC after six days normalized to respective control (TC6) **(d-f)**. *L. bicolor* transcriptome during confrontation with *T. harzianum* after three (LT3, MC) and six days (LT6, DC) normalized to respective single controls (LC3 and LC6) **(g,h)**.

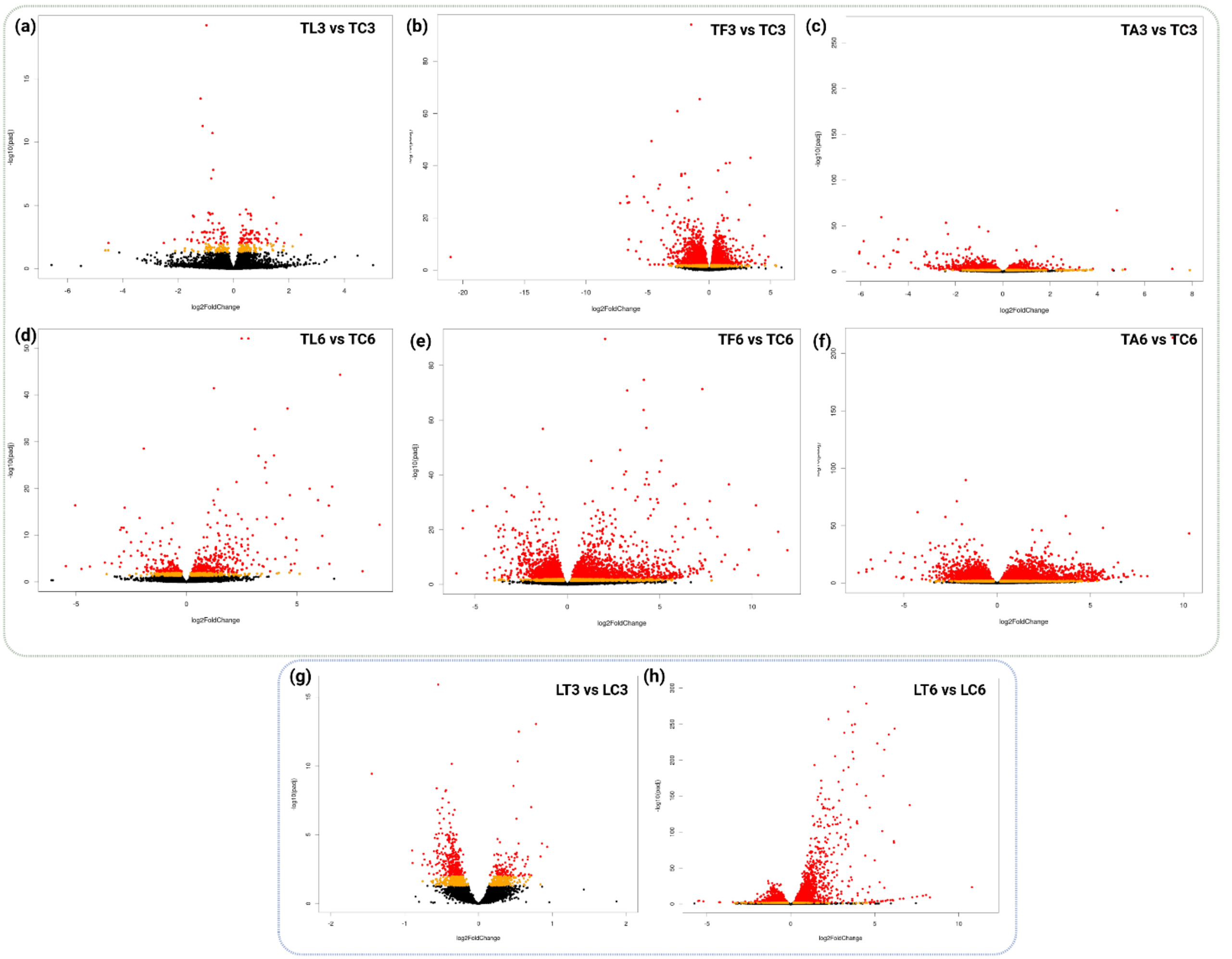

**Fig. S4:** Expression graphs and gene names based on SOTA cluster organization of the 57 common DEGs in *T. harzianum* after three days during confrontation with *A. alternata* (TA3), *F. graminearum* (TF3) and *L. bicolor* (TL3) of cluster 1 **(a)** and cluster 2 **(b).**

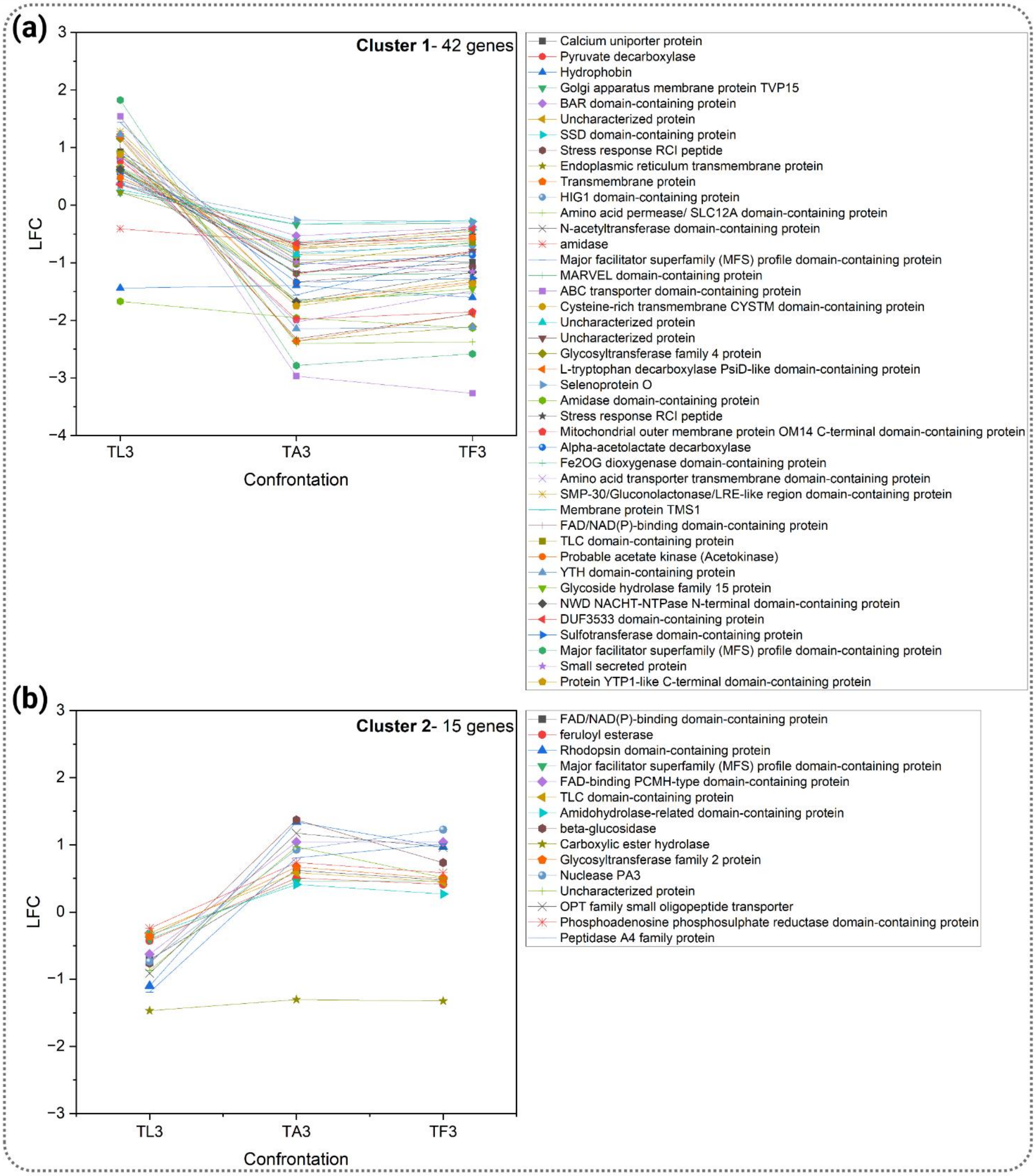

**Table S1** Genes identified as differentially expressed in *T. harzianum* and *L. bicolor* during plate confrontation compared to the single control conditions using DESeq2 analysis. Genes were assumed to be differentially expressed with an FDR < 0.01. Tables are showing the Gene IDs, UniProt entry, predicted protein names, length, baseMean, log2FoldChange (blue log2FC < 0, red log2FC > 0) and its SE as well as *p*-values and adj. *p*-values.

**Table S1-1:** DEGs identified in *T. harzianum* during interaction with *L. bicolor* during MC (three days).
**Table S1-2:** DEGs identified in *T. harzianum* during interaction with *A. alternata* during MC (three days).
**Table S1-3:** DEGs identified in *T. harzianum* during interaction with *F. graminearum* during MC (three days).
**Table S1-4:** DEGs identified in *T. harzianum* during interaction with *L. bicolor* during DC (six days).
**Table S1-5:** DEGs identified in *T. harzianum* during interaction with *A. alternata* during DC (six days).
**Table S1-6:** DEGs identified in *T. harzianum* during interaction with *F. graminearum* during DC (six days).
**Table S1-7:** DEGs identified in *L. bicolor* during interaction with *T. harzianum* during MC (three days).
**Table S1-8:** DEGs identified in *L. bicolor* during interaction with *T. harzianum* during DC (six days).

**Table S2** Common DEGs in *T. harzianum* during MC with *L. bicolor*, *A. alternata* and *F. graminearum* clustered using SOTA based on expression values. Displayed are the assigned clusters and log2FC (blue log2FC < 0, red log2FC > 0), UniProt entry, protein names, gene IDs, length, assigned GO terms and signal peptide prediction (SignalP).

**Table S3** Venny groups of common up- and downregulated DEGs in *T. harzianum*.
**Table S3-1** Downregulated DEGs in *T. harzianum* during MC (three days).
**Table S3-2** Upregulated DEGs in *T. harzianum* during MC (three days).
**Table S3-3** Downregulated DEGs in *T. harzianum* during DC (six days).
**Table S3-4** Upregulated DEGs in *T. harzianum* during DC (six days).

**Table S4** Gene ontology enrichment analysis results.
**Table S4-1:** Gene ontology (GO) enrichment analysis of upregulated DEGs in *L. bicolor* during confrontation with *T. harzianum* during MC (three days).
**Table S4-2:** FunCat enrichment analysis of upregulated DEGs in *L. bicolor* in confrontation with *T. harzianum* during MC (three days).

**Table S4-3:** KEGG enrichment analysis of upregulated DEGs in *L. bicolor* in confrontation with *T. harzianum* during MC (three days).

**Table S4-4:** Gene ontology (GO) enrichment analysis of upregulated DEGs in *L. bicolor* during confrontation with *T. harzianum* during DC (six days).

**Table S4-5:** FunCat enrichment analysis of upregulated DEGs in *L. bicolor* in confrontation with *T. harzianum* during DC (six days).

**Table S4-6:** KEGG enrichment analysis of upregulated DEGs in *L. bicolor* in confrontation with *T. harzianum* during DC (six days).

**Table S4-7:** Gene ontology (GO) enrichment analysis of common and unique upregulated DEGs in *T. harzianum* during confrontation with *A. alternata* and *F. graminearum* during MC (three days).

**Table S4-8:** Gene ontology (GO) enrichment analysis of unique upregulated DEGs in *T. harzianum* during confrontation with *L. bicolor* during MC (three days).

**Table S4-9:** Gene ontology (GO) enrichment analysis of common and unique upregulated DEGs in *T. harzianum* during confrontation with *A. alternata* and *F. graminearum* during DC (six days). .

**Table S4-10:** Gene ontology (GO) enrichment analysis of unique upregulated DEGs in *T. harzianum* during confrontation with *L. bicolor* during DC (six days).

**Methods S1** Cultivation of *P. x canescens*

Poplar micro-cuttings were transferred to SH medium to induce rooting and after four weeks plants with similar height and root length were selected and transferred into autoclaved jars (RR80, J. WECK GmbH & Co KG, Wehr-Öflingen, Germany) filled with 200 ml substrate (60 % vermiculite (1904, Jungepflanzen, Forchheim, Germany), 20 % fine sand (0.71-1.25 mm particle size, MGS Shopping, Hohenthann, Germany), 20 % perlite (KPP, Knauf, Iphofen, Germany)). The jars were sealed with a transparent gas and water permeable membrane (Z380059, Breathe-Easy®, Sigma, Deisenhofen, Germany).

**Fig. Methods S1:** Picture of *P. x canescens* and *T. harzianum* and *T. atrobrunneum* in jar cultivation system.

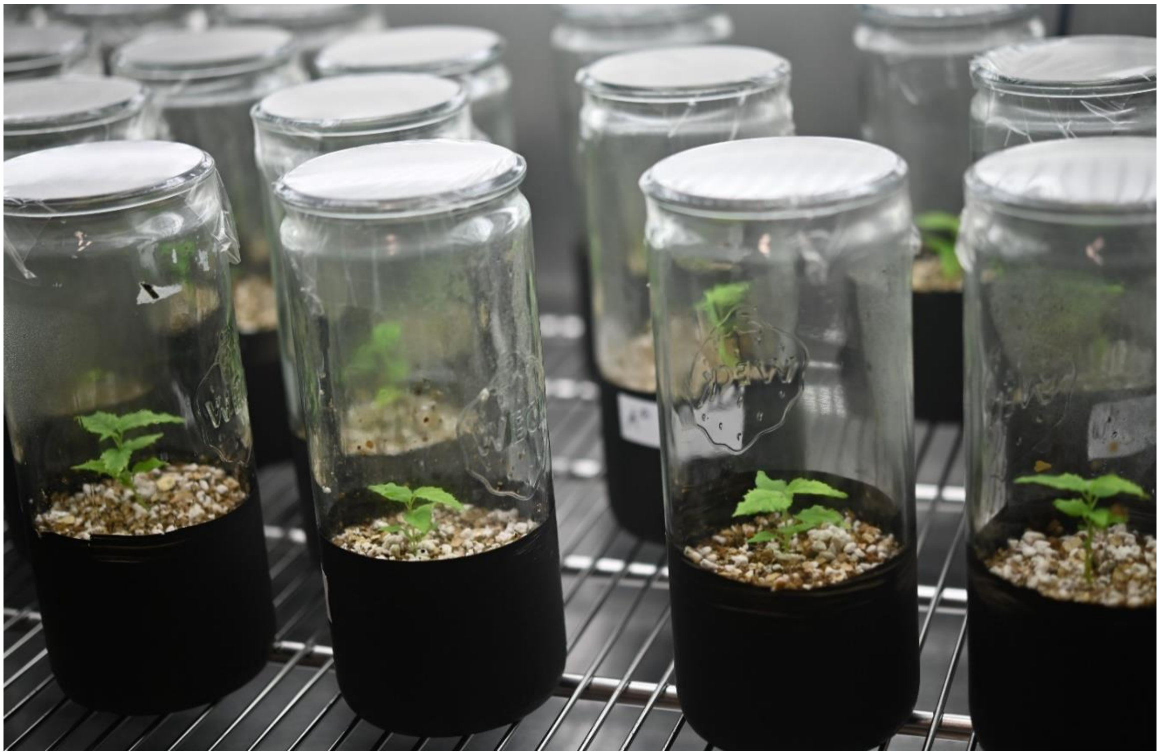

**Methods S2** Preparation and setup of olfactometer "race tube"-like system.

The olfactometer "race tube"-like system (Supplementary Fig. **S2a**) was composed of two 220 ml sample cups (391-0023, VWR, Darmstadt, Germany) and a 50 ml serological pipette (612-3696, VWR, Darmstadt, Germany). Holes were drilled into the side of the sample cups with a Forstner drill (ø = 16 mm, 2468320, kwb Germany GmbH, Stuhr, Germany) and both ends of the serological pipette were cut off with a hot scalpel under sterile conditions. Two sample cups and one serological pipette were assembled by connecting the two cups with the serological pipette and sealing the joints with physiologically harmless silicon (AquaSil, transparent, JBL, Neuhofen, Germany). The system was then filled with agar-based MMN media in a way that the media level was the same in both tubes and the horizontal serological pipette was filled in half, allowing media and air contact between both test tubes and a flat media surface (Supplementary Fig. **S2b**). To enable inoculation of *Trichoderma*, a hot needle was used to melt a small hole into in the middle of the serological pipette. Both sample cups and the melted hole were sealed with gas-permeable membrane (BR701364, Sigma, St. Louis, USA).

**Methods S3** RNA-extraction from fungal confrontations for transcriptomic analysis.

For RNA extraction fungal biomass was scraped with a sterile spatula from the interaction zone and immediately ground in liquid nitrogen and stored at -80°C till further processing. RNA was extracted from 4 biological replicates using the kit NucleoSpin RNA Plant and Fungi (Machery-Nagel, Düren, Germany) following the manufacturer's instructions. Integrity and total amount of RNA was detected by bioanalyzer and Qubit (RNA High Sensitivity assay, Aligent Technologies, Santa Clara, USA).
